# RNA binding of Hfq monomers promotes RelA-mediated hexamerization in a limiting Hfq environment

**DOI:** 10.1101/2020.08.11.244277

**Authors:** Pallabi Basu, Maya Elgrably-Weiss, Fouad Hassouna, Manoj Kumar, Reuven Wiener, Shoshy Altuvia

## Abstract

The RNA chaperone Hfq acting as a hexamer, is a known mediator of post-transcriptional regulation expediting basepairing between small RNAs (sRNAs) and their target mRNAs. However, the intricate details associated with Hfq-RNA biogenesis are still unclear. Previously, we reported that the stringent response regulator, RelA is a functional partner of Hfq that facilitates Hfq-mediated sRNA-mRNA regulation *in vivo* and induces Hfq hexamerization *in vitro*. Here, for the first time we show that RelA-mediated Hfq hexamerization requires an initial binding of RNA, preferably sRNA to Hfq monomers. By interacting with a Shine-Dalgarno-like sequence (GGAG) in the sRNA, RelA stabilizes the initially unstable complex of RNA bound-Hfq monomer, enabling the attachment of more Hfq subunits to form a functional hexamer. Overall, our study showing that RNA binding to Hfq monomers is at the heart of RelA-mediated Hfq hexamerization, challenges the previous concept that only Hfq hexamers can bind RNA.

## Introduction

As a rule, most RNA based regulation involves the function of RNA binding proteins including Hfq, ProQ, cold shock proteins and proteins of the CsrA family (*1–8*). Out of which, Hfq and its associated small regulatory RNAs were acknowledged as significant key players of a large network of post-transcriptional control of gene expression in Gram-negative bacteria. Acting as an RNA chaperone, Hfq facilitates basepairing between small regulatory RNAs and their target mRNAs, thereby leading to altered stability and/or translation of the target genes (*9–12*). The importance of Hfq for global RNA regulation has been substantiated through studies showing that Hfq interacts with a great number of different sRNAs and mRNAs species (*1, 2, 13*). Hfq was also shown to bind rRNAs and tRNAs, suggesting an effect on ribosome biogenesis and translation efficiency implicating Hfq as a global regulator (*14*).

Hfq structural studies showed that the protein forms a doughnut shaped homo-hexamer. The hexameric ring reveals four sites that can interact with RNA: proximal and rim faces interacts with uridines present in the 3’ end of sRNAs, distal face interacts with ARN motifs present in the target mRNAs and C-terminal tail ensures the release of the RNAs from Hfq, enabling Hfq recycling (*15–17*). In addition to its affinity for RNA, Hfq interacts with components of the RNA decay machinery such as poly(A) polymerase, polynucleotide phosphorylase, RNase E and the transcription termination factor Rho (*18*-*22*).

*In-vitro* studies indicated that Hfq transitions from monomer to hexamer at about 1µM of Hfq protein and that RNA-bound Hfq hexamer is a stable complex (*23*). At higher concentrations, Hfq predominantly forms multimers, whereas upon dilution, the subunits dissociate, indicating that multimerization depends on the Hfq microenvironment and that the interactions are reversible (*23*). Mutations in Hfq that impair RNA binding either strongly destabilize the hexamer or prevent hexamer association to multimers, indicating that RNA binding is coupled to hexamer assembly (*24–26*). Whether RNA binding coincides with hexamerization which requires initial disassembly of Hfq, assuming that RNA can bind individual Hfq subunits to form a new RNA-bound complex or whether hexamers are the only forms capable of RNA binding which necessitates random recycling of new RNAs on the surface of Hfq are some of the unresolved issues regarding the Hfq-RNA biogenesis. Both the options also raise the possibility that other regulators chaperones Hfq-RNA biogenesis.

While investigating expression regulation by RyhB sRNA, we discovered that the stringent response regulator protein RelA is a functional partner of Hfq mediating RyhB-target regulation (*27*). We suggested that RelA impacts RyhB-target mRNA regulation by promoting assembly of Hfq monomers into hexamers and thereby enabling low and ineffective concentrations of Hfq to bind RNA (*27*).

The RelA protein of *Escherichia coli* is a ribosome-dependent (p)ppGpp synthetase that is activated under conditions of amino acid starvation (*28, 29*). Once produced, (p)ppGpp modifies the activities of multiple cellular targets, including enzymes for DNA replication, transcription, translation, ribosome assembly, cellular metabolism and genome stability (*28, 30–32*). RelA synthetase activity resides within the amino terminus of the protein whereas the carboxy terminus enables regulation of the synthetase function in a ribosome-dependent manner (*33, 34*).

Here we show RNA binds Hfq monomers and that RelA by interacting with a specific sequence in the sRNA, stabilizes the initially unstable complex of RNA-Hfq monomer, promoting the association of additional Hfq subunits to form the hexameric complex. Overall, our study challenges the previous concept that only Hfq hexamer can bind RNA and introduces a new chaperone-like regulator that mediates RNA-bound Hfq hexamerization.

## Results

### RelA amino terminus facilitates repression of RyhB targets by RyhB

To identify domains in RelA that promote Hfq activity, we carried out deletion mapping in which either the N-terminus or the C-terminus of RelA were eliminated. The genetic system used to test these constructs included RyhB target reporters (*sdhC*-*lacZ* and *sodA*-*lacZ*) as single copies, chromosomally encoded Hfq and plasmids encoding RelA and RyhB in Δ*relA*Δ*ryhB* strain. RelA and truncated RelA carrying only the N-terminal domain (pRelA-ΔCTD) enabled repression of RyhB targets by RyhB (Figure 1A and Figure S1). In contrast, RelA-ΔNTD, carrying only the C-terminal domain of RelA failed to enable repression, indicating that the N-terminal domain of RelA is essential for RyhB-mediated regulation of *sdhC* and *sodA*. We used the same genetic system to isolate RelA mutants by subjecting the RelA gene to random mutagenesis. Two single point mutations in RelA, C289Y and T298I, that reduced the repression of RyhB target genes were clustered in one helix of the RelA N-terminal domain (Figure 1A, Figure S1, Figure S2).

**Figure 1.**
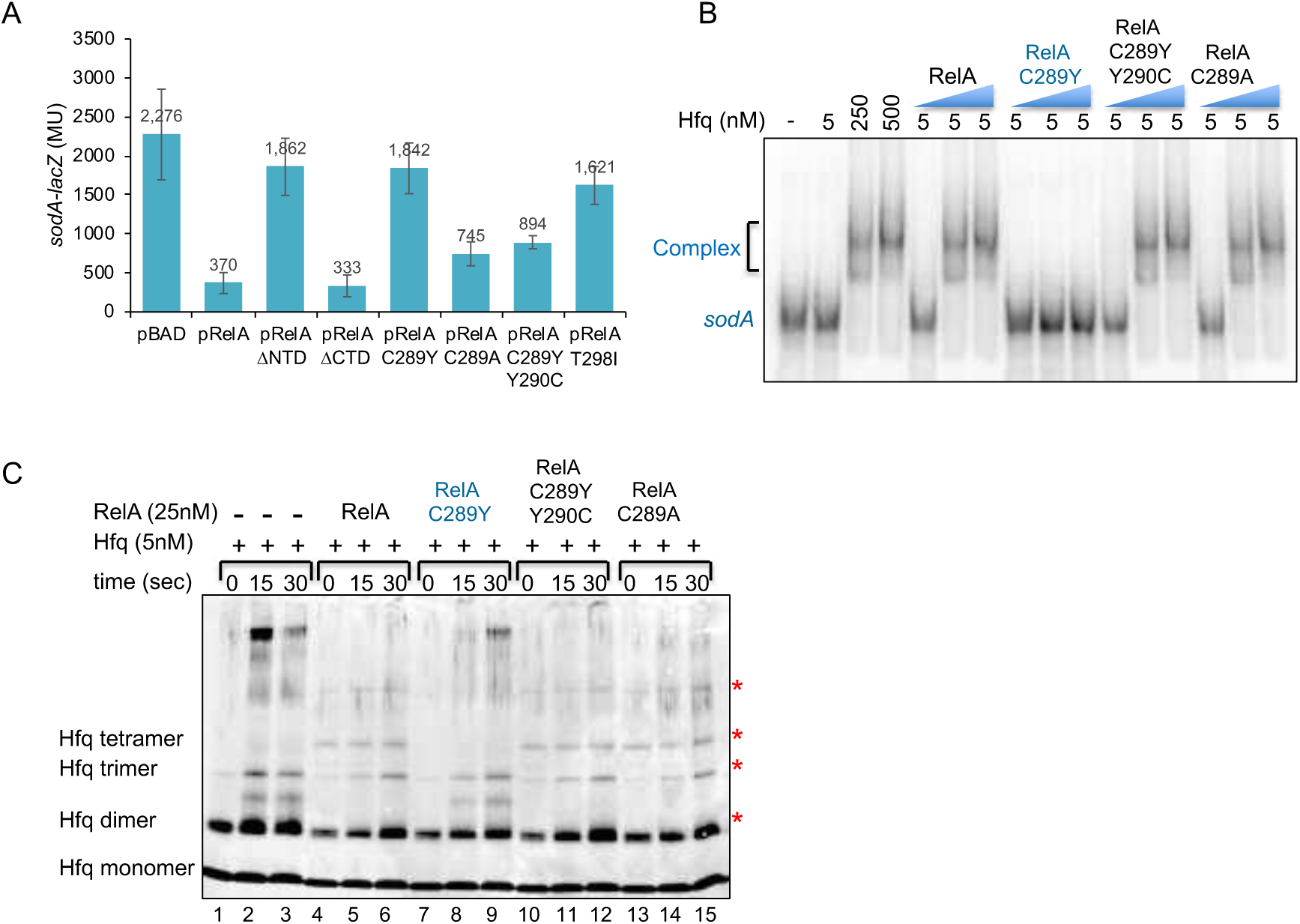
RelA amino terminus domain induces Hfq assembly. (**A**) β-galactosidase assay to determine the effect of plasmids encoded RelA alleles on repression of *sodA-lacZ* target gene fusion by RyhB. Expression of RelA from BAD promoter was induced with (0.2%) arabinose. (**B**) RelA facilitates binding of RNA to Hfq *in vitro* (EMSA). Gel mobility shift assay of radiolabeled *sodA* RNA incubated with Hfq without and with 50 nM, 250 nM and 500 nM of purified wild type or RelA mutant proteins as indicated (blue triangle). Incubations were carried out at 22°C for 10 min and the products were separated by 4% native gel electrophoresis. Unbound (*sodA*) and bound (complex) RNA is indicated in the figure. (**C**) RelA enhances the multimerization of Hfq protein (western). Hfq was incubated with or without purified RelA proteins for 10 min at 22°C. Thereafter, the products were cross-linked with 0.2% of glutaraldehyde at 22°C. Samples were collected at the indicated time points and the reactions were stopped with 200 mM fresh glycine. The proteins separated in 4-20% MOPS gradient gels were detected using *α* Hfq. Red asterisk indicates the formation of new multimers.

To test whether RelA mediated regulation is sequence-specific or stemmed from structural elements; we changed the cysteine residue (small and non-polar) at position 289 to alanine harboring characteristics similar to cysteine. RelA:C289A rescued 50% of *sodA-lacZ* repression and 85% of *sdhC-lacZ* repression by RyhB (Figure 1A, Figure S1). As wild type RelA carries tyrosine, an aromatic amino acid residue at position 290, the mutational change C289Y resulted in two consecutive tyrosine residues that were expected to cause steric hindrance because of their bulky side chains (*35*). The double mutant RelA:C289Y;Y290C in which the tyrosine residue at position 290 was changed to cysteine was more effective in mediating repression than the single C289Y mutant, suggesting that C289Y causes steric hindrance and that RelA-mediated regulation relies primarily on structural elements (Figure 1A, Figure S1).

As RelA mutants affecting repression of RyhB targets reside in the amino terminus of RelA, we examined whether (p)ppGpp production correlated with the RelA regulatory activity of basepairing RNAs. The double mutant strain Δ*relA*Δ*spoT* fails to grow in M9 minimal medium unless supplemented with a plasmid producing (p)ppGpp. *ΔrelA*Δ*spoT* cells carrying the empty vector plasmid and pRelA-ΔNTD did not grow on minimal plates, whereas the growth of RelA mutants unable to facilitate RyhB target repression (pRelA:C289Y, pRelA:T298I) and those supporting repression by RyhB (pRelA:C289A, pRelA:C289Y;Y290C) was comparable to cells expressing wild type RelA, indicative of (p)ppGpp production (Figure S3). The analysis of RelA:Q264E, a (p)ppGpp synthetase deficient RelA mutant (*36*) that enabled *sdhC-lacZ* repression by RyhB further confirmed that (p)ppGpp production was not associated with RelA regulation of basepairing RNAs (Figure S1 Figure S3), indicating that (p)ppGpp production and RelA-mediated repression regulation are distinct functions, although both reside in the N-terminal domain.

### RelA amino terminus induces Hfq assembly

Previously, we have shown that purified wild type RelA enhanced the RNA binding activity of Hfq and Hfq oligomerization *in vitro* (*27*). The *in vivo* phenotype of RelA mutants prompted us to examine their effect on Hfq RNA binding and on Hfq quaternary structure. Gel mobility shift assays showed that low concentrations of Hfq (5 nM) were insufficient to bind *sodA* RNA unless incubated in the presence of RelA (Figure 1B). The *in vivo* inactive mutant RelA:C289Y failed to facilitate binding of RNA by Hfq, whereas the active suppressor mutants; RelA:C289A and RelA:C289Y;Y290C enhanced the binding activity of Hfq similar to wild type RelA (Figure 1B).

To evaluate the effect of RelA mutants on Hfq quaternary structure, Hfq protein incubated with RyhB sRNA and with or without RelA was exposed to glutaraldehyde, a protein crosslinking reagent. The reaction products were separated by SDS-PAGE and detected using *α* Hfq antibody. The pattern of Hfq oligomerization obtained upon incubation with RelA:C289A and RelA:C289Y;Y290C was similar to that detected with wild type RelA (note the presence of tetramers), whereas the pattern of Hfq oligomerization obtained with RelA:C289Y was similar to the pattern detected with Hfq alone (Figure 1C). Combining the *in vivo* and the *in vitro* results indicates that RelA supports expression regulation by basepairing RNAs and enhances Hfq RNA binding by facilitating oligomerization of Hfq. Furthermore, the function that supports basepairing RNAs is distinct from the synthetase activity although both reside in RelA amino terminus.

### RelA binds RNA bound by Hfq

The functional interaction between RelA and Hfq motivated us to investigate the *in vivo* molecular interaction between these two proteins. Co-immunoprecipitation using *α*RelA showed that Hfq precipitates in a complex with RelA and identified RNA as the mediator connecting between RelA and Hfq (Figure 2A). Hfq did not precipitate with RelA when the lysate was treated with RNase A, suggesting that in the absence of RNA the complex disassembles. Likewise, Hfq did not precipitate with RelA:C289Y mutant that is unable to promote RNA binding by low concentrations of Hfq or induce Hfq assembly. These results demonstrate that *in vivo*, RNA links between RelA and Hfq forming a complex.

**Figure 2.**
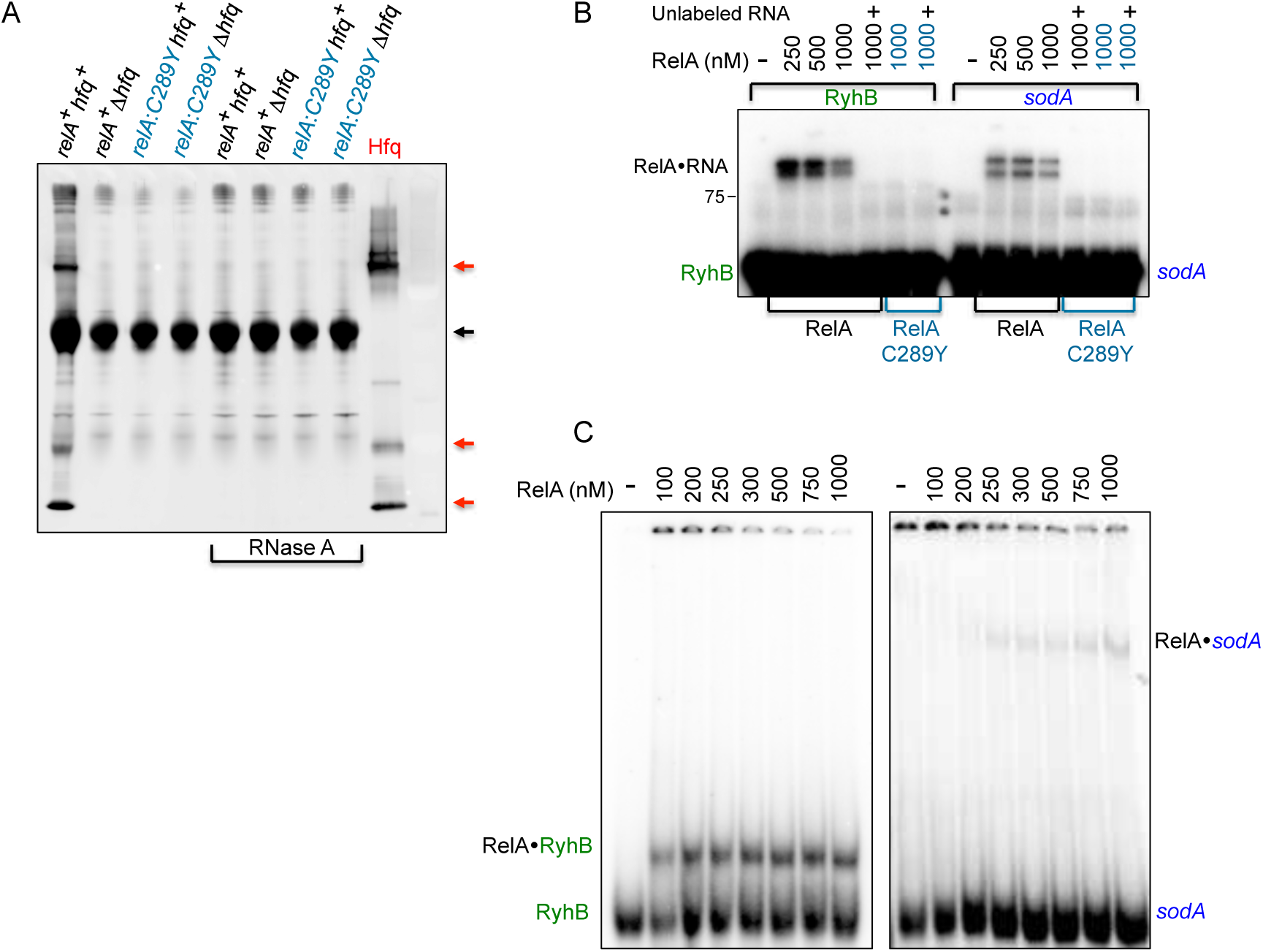
RelA binds RNA. (**A**) Co-immunoprecipitation carried out with cell lysates of wild type RelA (black) and RelA C289Y mutant (blue) as indicated. The lysates were treated with RNase A (100 μg/ml) or left untreated. Pull down was carried out with *α* RelA followed by incubation with Protein A sepharose beads. Hfq was detected by western using *α* Hfq antibody. Purified Hfq (100 nM) was used as control. Red arrows indicate different forms of Hfq, while black arrow indicates the heavy chain of *α* RelA antibody. (**B**) *In vitro* binding of RyhB and *sodA* by RelA. Wild type RelA or RelA:C289Y incubated with labeled RNAs (1 nM) were UV cross-linked. Competitor unlabeled RNA (100 nM) was added to the reaction mixtures as indicated. The binding products were analyzed by 15% SDS-PAGE. The estimated MW of the RNA•RelA complex is 107 kDa to 113 kDa. (**C**) Gel mobility shift assay of radiolabeled *sodA* (56 nt) or RyhB (50 nt) RNAs incubated with increasing concentrations of RelA as indicated. Incubations were carried out at 22°C for 10 min. The products were UV cross-linked before loading on 4% native gel electrophoresis.

To visualize direct binding between RelA and RNA, *in vitro* labeled *sodA* or RyhB RNAs were incubated with purified wild type RelA and RelA:C289Y mutant followed by UV cross-linking. Thereafter, the unbound and thus unprotected RNA residues were subjected to degradation by RNase A or left intact. Proteins covalently bound to untrimmed, labeled RNA (Figure 2B) or to truncated RNA residues (Figure S4) were then detected in SDS gels. The results demonstrate that wild type RelA binds both RNAs however RyhB binding by RelA is much stronger when compared to RelA binding affinity for *sodA*. The addition of unlabeled competitor RNAs eliminated binding of the labeled RNAs and RelA:C289Y showed no binding, indicating RelA:C289Y that is unable to induce Hfq assembly is also incapable of RNA binding (Figure 2B and Figure S4). As for the decrease in binding detected with 1μM of RelA, we suspect it is due to the formation of higher molecular weight complexes that failed to enter the gel.

The higher affinity of RelA for RyhB was further confirmed by gel mobility shift experiments. EMSA presented in Figure 2C shows that ∼100 nM of RelA binds approximately 50% of RyhB sRNA, whereas the binding affinity of RelA to *sodA* mRNA is much weaker. Unlike Hfq which when at high concentrations binds most of the RNA, the increase in RNA binding by increasing concentrations of RelA is much more moderate. As the *in vivo* complex of RNA bound by Hfq and RelA is sufficiently stable to be precipitated by *α* RelA antibody, yet *in vitro* RNA binding by RelA is limited, we suspect that RelA binding of RNA that is structurally modified by Hfq is more efficient. In the absence of the Hfq RNA pairing, the interaction of RelA with unaltered RNA is more elusive.

### RelA binds RNA with a specific sequence

To define domains in *sodA* and RyhB RNA that interact with RelA, we mapped the sites protected by RelA using dimethyl sulfate (DMS) that methylates unpaired adenosine and cytidine residues or RNase T1 that is specific for unpaired guanosine residues. The modified nucleotides and cleavage sites were mapped by primer extension. The results displayed in Figure 3A-C and summarized in Figure 3D show that RelA protects the sequence GGAGA in both *sodA* and RyhB. RyhB also consists of a variation of this sequence (GGAAGA) but RelA did not protect this site. The pattern of RNA probing upon incubation with RelA:C289Y mutant was similar to the pattern obtained in the absence of RelA, further confirming that C289Y mutant does not bind RNA. To confirm that GGAGA is the site RelA interacts with, we changed this sequence to ACUCU in *sodA* (*sodAm*) (Figure 3ABD). The pattern of RNA probing of *sodAm* incubated with wild type RelA or RelA:C289Y mutant was identical to the pattern detected in the absence of RelA indicating that RelA binds the sequence GGAGA which intriguingly resembles the ribosome binding Shine-Dalgarno sequence (Figure 3ABD).

**Figure 3.**
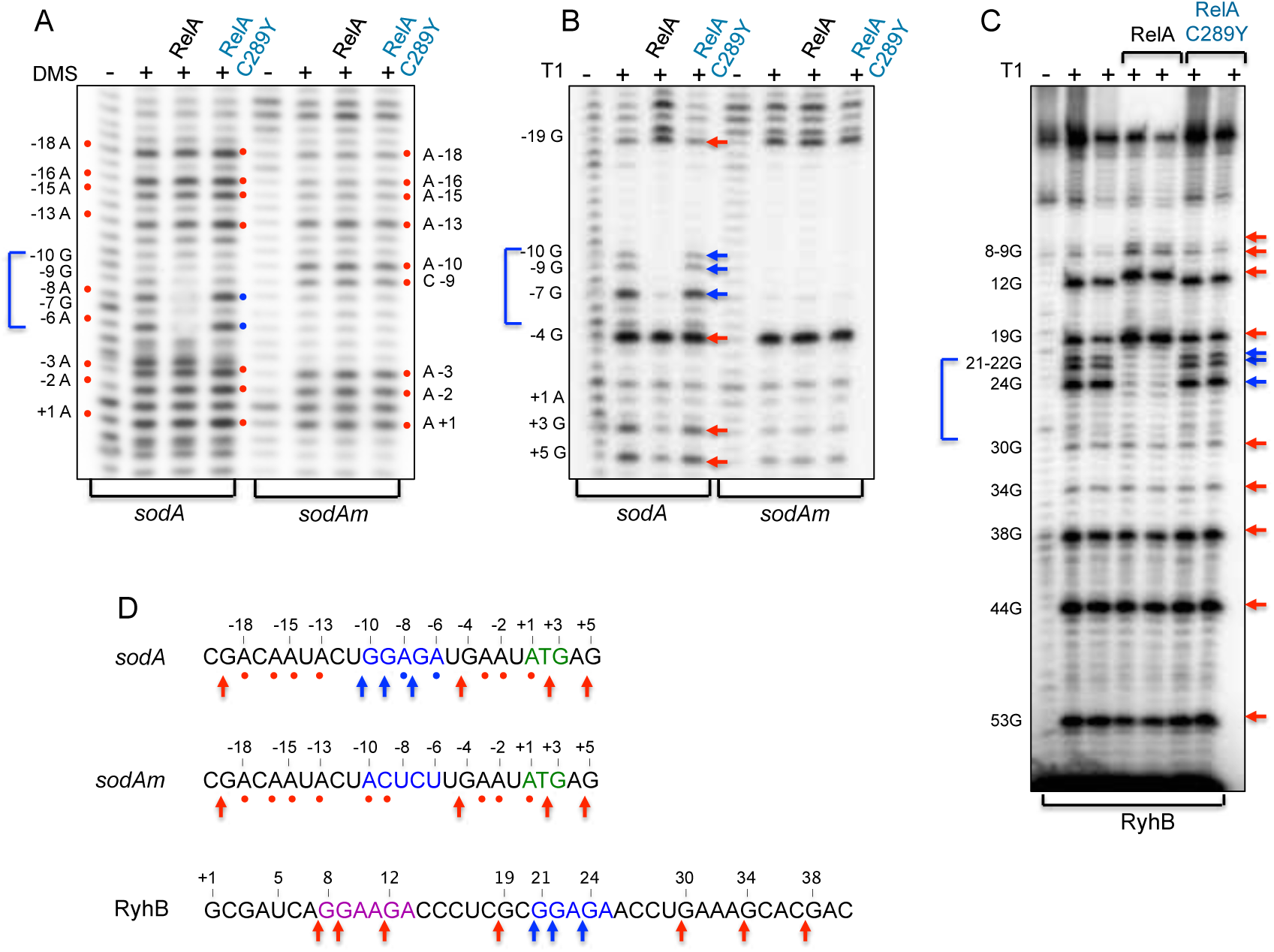
RelA interacts with RNA through a specific sequence. (**A**) Foot printing of RelA using DMS modification. Wild type (black) and mutant (blue) RelA proteins (5 pmol) incubated with (0.5 pmol) RNAs were exposed to DMS modification (0.3%) for 5 min at 25°C. *sodA* carries an intact GGAGA sequence while *sodA*m carries ACUCU. Reverse transcription of untreated (-) and DMS treated (+) RNA samples. The red circles indicate the positions methylated by DMS. The numbers on the right and left indicate the sequence position relative to the nucleotide A of the start codon of *sodA* (+1). Wild type RelA protects residues A-8 and A-6 from methylation (blue circles). Nucleotides A-10 and C-9 in *sodA*m RNA are methylated (red circles) in the presence of either wild-type or RelA:C289Y mutant. (**B, C**). Foot printing of RelA using RNase T1. RNAs and proteins incubated as in A were treated by RNase T1 (0.1 U) for 5 min at 37°C (B) or with 0.2 U and 0.4 U (C). Reverse transcription of untreated (-) and RNase T1 treated (+) RNA samples. The numbers on the left indicate the sequence position relative to the nucleotide A of the start codon of *sodA* and the transcription start site of RyhB (+1). The red arrows indicate the positions of the G residues cleaved by RNase T1, while the blue arrows represent the regions of protection. (**D**) RelA protects GGAGA sequence of RyhB and *sodA*. The sequences GGAGA (blue), AUG (green) and variant GGAAGA (purple) are denoted. In *sodA*m GGAGA was changed to ACUCU. Red circles and arrows indicate strong modification and cleavage sites. Blue circles and arrows indicate the region protected by RelA. The products were analyzed in 6% acrylamide 8M urea sequencing gel.

### RelA induces Hfq assembly by binding RNA with GGAGA

To further confirm that RelA mediated Hfq assembly requires interaction with GGAGA, we investigated the assembly pattern of Hfq in the presence of wild type and mutants of *sodA* and RyhB RNAs. To this end, we constructed *sodA* that lacks the GGAGA region (*Δ*SD) and RyhB in which GGAGA was changed to CAUCU (RyhBm). In RyhBm, the variant site GGAAGA was also mutated to GGUUCA (Figure S5). Protein crosslinking of Hfq showed that in the absence of RelA, the addition of any of the RNAs had no effect on assembly of low concentrations of Hfq (Figures 4A and C lanes 1-9). In the presence of RelA, the addition of wild type *sodA* or RyhB resulted in an increase in Hfq oligomerization (Figures 4B and D compare lanes 1-3 to 4-6), whereas the assembly pattern of Hfq presented with *sodA*-*Δ*SD or with RyhBm was similar to that detected without any RNA (Figures 4B and D lanes 1-3 and 7-9). Taken together, the results indicate that in the absence of RelA, RNA has no effect on oligomerization of low concentrations of Hfq. However, RNA plays a significant role in RelA-induced Hfq hexamerization that is driven by RelA interacting with RNA carrying a GGAGA site.

**Figure 4.**
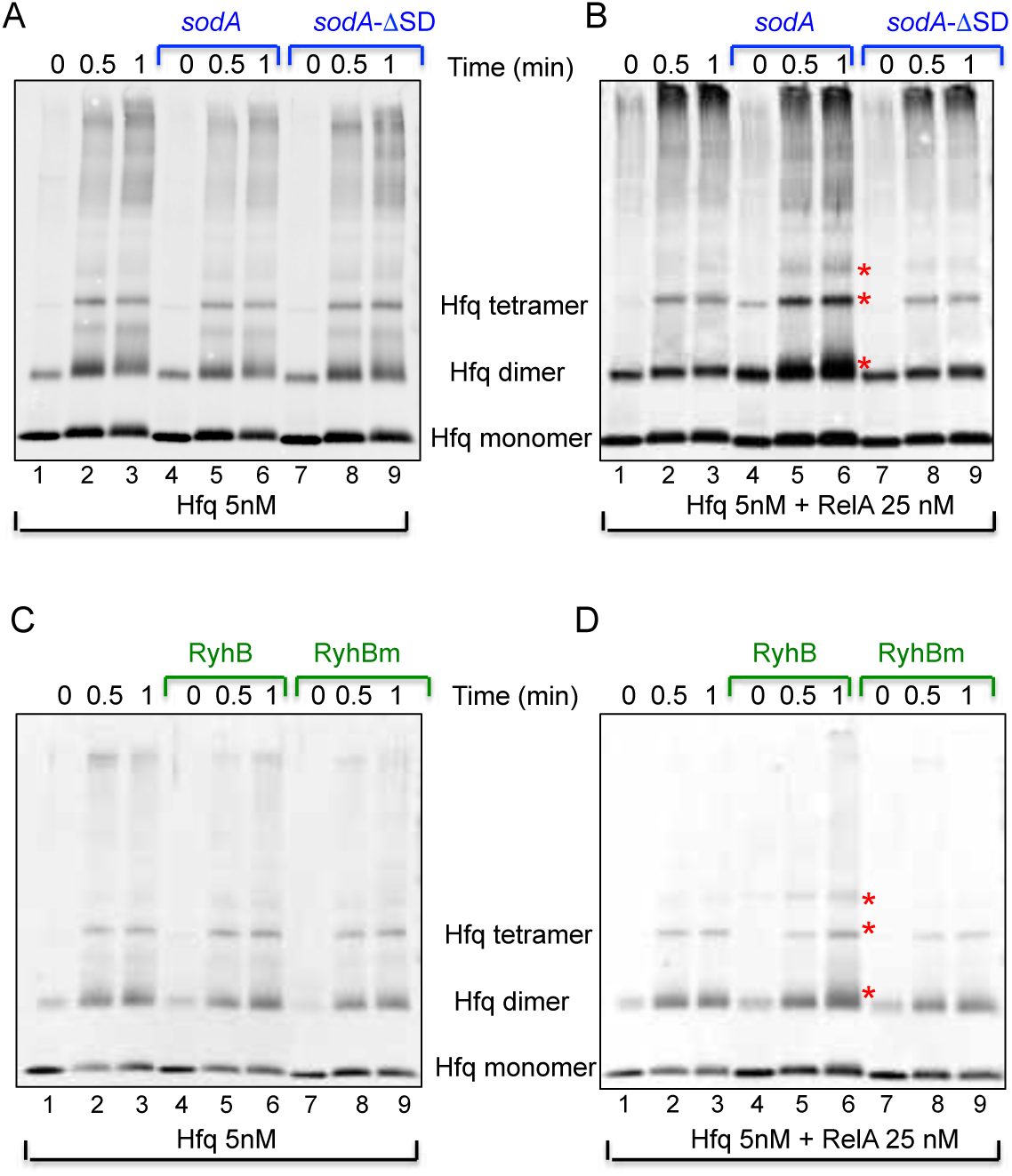
RelA mediated Hfq assembly requires interaction with GGAGA sequence (western using *α* Hfq antibody). (**A, C**) In the absence of RelA, RNA has no effect on Hfq multimerization. Reactions of Hfq incubated without or with RNA at 22°C for 10 min. were UV cross-linked followed by protein crosslinking with 0.2% of glutaraldehyde. The proteins separated in 4-20% MOPS gradient gels were detected using *α*Hfq. (**B, D**) RelA induces Hfq multimerization when presented with RNA carrying GGAGA. Reactions of Hfq incubated with RelA, without or with RNA including RyhB, RyhBm *sodA* and *sodA*-ΔSD were treated as in A. Asterisk indicates the formation of new Hfq multimers detected using wild type RNAs in the presence of RelA.

### RelA stabilizes complexes of RNA associated with Hfq monomers to form hexamers

To follow the steps of RelA induced Hfq assembly to hexamers, Hfq (5 nM) and labeled RyhB RNA, with or without RelA were crosslinked and the products were separated on SDS gels. Intriguingly, we detected binding of labeled RyhB to one Hfq monomer (Figure 5A). The binding was visible only in the presence of RelA, unlabeled RNA competed with the labeled one for Hfq binding and reactions carrying RelA:C289Y mutant showed no binding, indicating that RelA stabilizes complexes of RNA associated with Hfq monomers by interacting with the RNA. Similarly, incubation of labeled *sodA* with Hfq in the presence of RelA resulted in formation of *sodA*•Hfq complex. However, the complex *sodA*•Hfq was significantly weaker compared to RyhB•Hfq, suggesting that sRNA is a much better substrate for RelA (Figure S6). Incubation of labeled *sodAm* RNA with RelA and Hfq resulted in no binding, indicating the preference of RelA to RNA with GGAGA (Figure S6).

**Figure 5.**
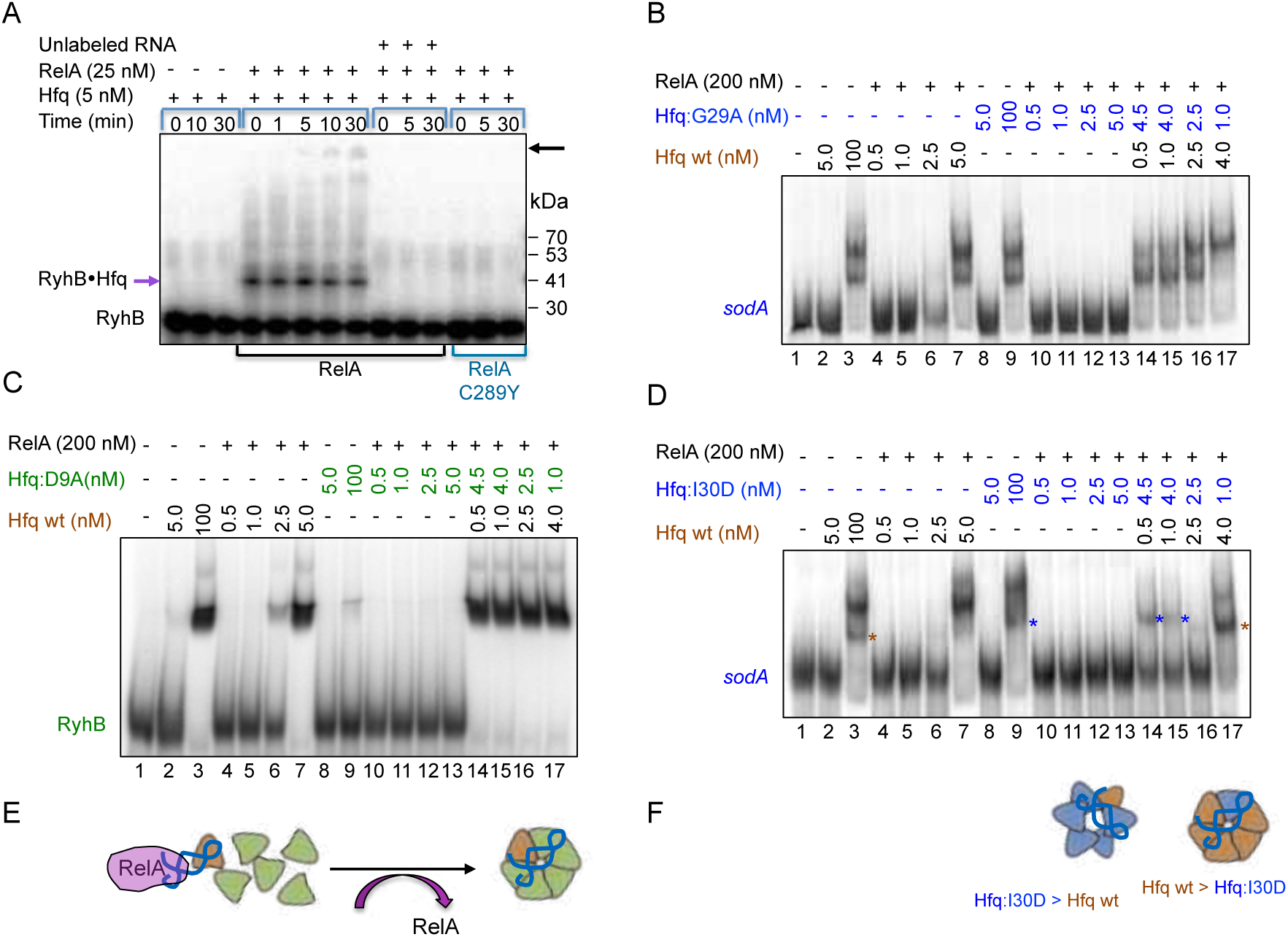
RelA stabilizes the binding of RNA to Hfq monomer and enables further assembly. (**A**) RelA facilitates RyhB binding to Hfq monomers. Reaction mixtures of Hfq incubated for 10 min at 22°C with labeled RyhB (1 nM) without or with RelA or RelA:C289Y were UV cross-linked followed by protein crosslinking with 0.2% glutaraldehyde. The cross linking was stopped with 200 mM of fresh glycine and the products analyzed in 4-20% MOPS gradient gel. Unlabeled competitor RNA (100 nM) was added where indicated. The estimated MW of the RNA•Hfq monomer complex (purple arrow on lower left side) is ∼ 40 kDa. Note that the addition of RNA alone to low levels of Hfq (5 nM) does not result in Hfq-RNA stable binding. Also, RelA:C289Y does not enable the binding of RNA to Hfq monomers. In the presence of RelA, as time of cross-linking progressed, higher forms of Hfq•RyhB emerged indicated by a black arrow (**B**) Gel mobility shift assay carried out with different ratios of Hfq to Hfq:G29A distal mutant (blue) incubated with labeled *sodA* distal RNA for 10 min at 22°C followed by 4% native gel electrophoresis. The binding of as low as 0.5 nM of Hfq to *sodA* RNA is necessary and sufficient to enable further assembly with Hfq:G29A subunits. See illustration of Hfq assembly pathway in E (**C**) As in B except that RyhB and a proximal face Hfq mutant Hfq:D9A were used. See illustration of Hfq assembly pathway in E (**D**) As in B, except that RNA binding to Hfq:I30D forms a complex that is different from that formed by wild type (see illustration of the two forms of hexamers in F. Brown asterisk denotes the position of the wild type complex (lane 3) whereas blue asterisk denotes the position of the Hfq:I30D complex (lane 9). The position of the complex in lanes 14 and 15 is shifted towards wild type as the ratio of wild type to I30D is increasing (lane 17). (E) Illustration of Hfq assembly induced by RelA. RelA stabilizes the binding of RNA to wild type Hfq monomer (brown) and enables the addition of mutant Hfq subunits (green). (F) Illustration of Hfq hexamers composed mainly by wild type (brown) or by mutated Hfq subunits (blue).

Our results suggest that RelA stabilizes an initial complex of RNA associated with Hfq monomer and thereby enables attachment of additional monomers to form Hfq hexamers. To show that preliminary RNA binding to Hfq monomers is necessary and sufficient for RelA to initiate Hfq assembly, we mixed limiting levels of wild type Hfq with comparatively higher levels of either Hfq distal mutant (I30D or G29A) and *sodA* (distal RNA unable to bind distal mutant) or with Hfq proximal face mutant (D9A) and RyhB (proximal RNA unable to bind proximal mutant). We based this experiment on the assumption that RelA stabilization of the complex formed by the binding of RNA to wild type Hfq monomers allows additional mutated Hfq subunits to join the initial complex to from hexamers. Figure 5BC shows that RelA fails to facilitate assembly of extremely low inactive levels (< 5 nM) of wild type Hfq when incubated along with *sodA* or RyhB RNA (lanes 4-6), all the more so of Hfq:G29A (distal) mutant incubated with *sodA* (Figure 5B lanes 10-13) and Hfq:D9A (proximal) incubated with RyhB (Figure 5C lanes 10-13). However, mixing labeled *sodA* RNA with low, inactive levels of wild type Hfq (0.5 nM) and 4.5 nM of distal face Hfq:G29A subunits that are unable to bind *sodA* resulted in *sodA* binding and heterogeneous complex formation (Figure 5B; lanes 14-17), indicating that the little RNA binding of *sodA* by wild type Hfq monomers enabled stabilization of the complex by RelA. The binary complex served as an anchor for further attachment of Hfq:G29A subunits leading to the formation of a mixed subunits hexamer (see illustration in Figure 5E). Likewise, mixing inactive levels of wild type Hfq with Hfq:D9A proximal face mutant that is unable to bind RyhB resulted in RyhB binding (Figure 5C; lanes 14-17) further confirming that initial RNA binding to wild type Hfq is necessary and sufficient to initiate hexamer formation by RelA.

The position of the complex formed by RNA binding to Hfq:I30D is slightly different from the one detected with wild type Hfq (Figure 5D; compare lanes 3 and 9) Interestingly, we noticed that the position of the complex obtained by mixing wild type (0.5 nM) with Hfq:I30D (4.5 nM) is similar to that detected with Hfq:I30D alone (Figure 5D; lanes 14,15). However, as the number of Hfq wild type subunits increases the position of the complex is shifted towards the wild type position (Figure 5D; lanes 16,17 and illustration in Figure 5F), further confirming that the hexamer is formed by mixing different subunits and as the ratio changes the complex’s position changes too. Together the results strongly demonstrate that RelA stabilization of the preliminary complex of Hfq subunit bound by RNA enables the attachment of additional subunits to form hexamers.

### (p)ppGpp synthesis and RNA binding are mutually exclusive functions of RelA

The observation that RelA:C289Y mutant failed to enable repression regulation of RyhB targets by RyhB yet it produced (p)ppGpp (Figure S3, Figure S7A) prompted us to investigate the interaction between these two functions. *In vivo* assays of (p)ppGpp production carried out with chromosomally encoded *relA*^+^ and *relA*:C289Y strains showed that upon amino acid starvation both strains produced similar levels of (p)ppGpp. In the presence of a plasmid expressing RyhB, (p)ppGpp production by wild type RelA decreased by 1.5-fold, whereas RelA:C289Y was much less affected by the RNA, indicating that RNA binding inhibits the synthetase activity of RelA and further confirming that RelA:C289Y is unable to bind RNA (Figure 6A)

**Figure 6.**
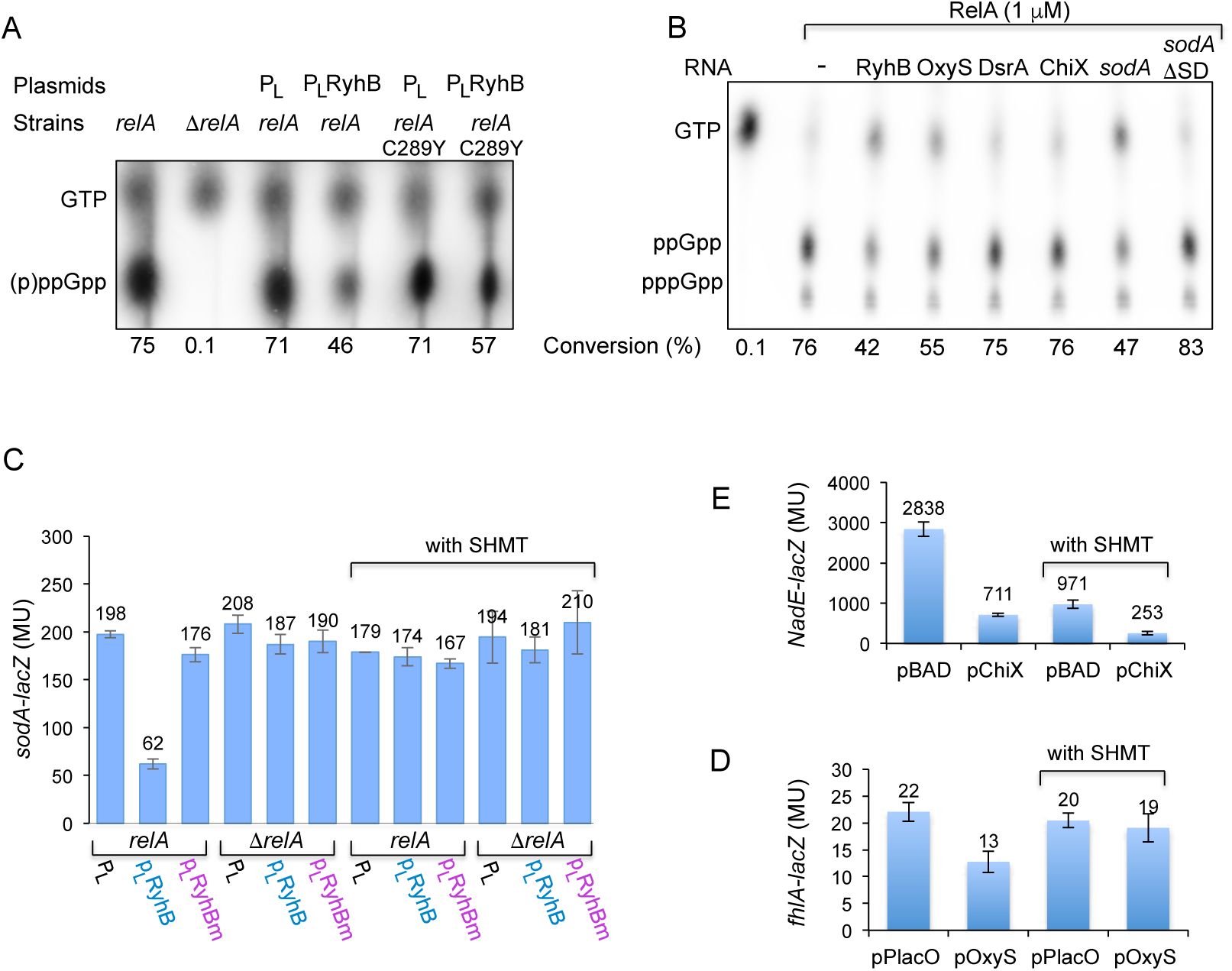
(p)ppGpp production and RNA binding are two mutually exclusive functions of the stringent response regulator RelA (thin layer chromatography). (**A**) *In-vivo* (p)ppGpp production is inhibited by RyhB. *E. coli* strains; *relA*^+^, Δ*relA*, and *rel:*C289Y (chromosomally encoded) carrying plasmids as indicated were assayed for (p)ppGpp production as described in material and methods. The intensity of the spots was determined by the Image Quant software and percentage of (p)ppGpp production of the total was calculated (% conversion). (**B**) *In-vitro* (p)ppGpp production is inhibited by specific RNAs. Purified RelA was incubated with either RyhB, OxyS, DsrA, ChiX, *sodA* or *sodA*-ΔSD and assayed by TLC. The intensity of the spots was determined by the Image Quant software and percentage of (p)ppGpp production of the total was calculated (% conversion). DsrA and ChiX lack a GGAGA site (see Figure S5). (C,D,E) RelA mediated basepairing regulation under normal growth conditions and in response to amino acid starvation with serine hydroxamate (with SHMT). β-galactosidase assays of target fusions in the presence of their corresponding sRNAs (RyhB/*sodA*, ChiX/*nadE* or OxyS/*fhlA*). ChiX expression was induced by 0.2% arabinose from the BAD promoter. Constitutive expression of plasmid encoded RyhB and OxyS in Δ*ryhB* and wild type respectively.

The observation that RyhB affected the synthetase activity, prompted us to examine whether RelA is specific to RyhB and its targets. *In vitro* (p)ppGpp assay carried out with RelA incubated with OxyS sRNA carrying GGAG or with DsrA and ChiX sRNAs that lack this specific sequence (Figure S5) showed that RNAs with GGAG (RyhB, OxyS and *sodA*) decreased production of (p)ppGpp by RelA. In contrast, DsrA, ChiX and *sodA*-ΔSD had no effect on (p)ppGpp production (Figure 6B). Furthermore, (p)ppGpp production in the presence of *sodA* in which the GGAGA sequence was mutated to ACUCU was unaffected (Figure S7). Taken together the results demonstrate that RelA binds RNAs with GGAG and this binding interferes with its synthetic activity.

### sRNAs with GGAG trigger RelA function *in vivo*

Our results suggest that conditions of amino acid starvation will inhibit RelA regulatory activity of basepairing RNAs *in vivo*. *β*-galactosidase assays of *sdhC-lacZ* or *sodA-lacZ* carried out in the presence of serine hydroxamate showed that upon starvation, RelA mediated repression regulation by RyhB was impaired (Figure 6C). Given that RyhB and both its targets *sdhC* and *sodA* carry GGAG (Figure S5), RelA regulation of basepairing RNAs could be due to RelA binding of either RyhB or its targets or both. To examine which of the RNAs triggers RelA regulation we mutated the GGAG RelA binding site in RyhB. To make sure that the core sequence that is responsible for RyhB regulation of its targets remained intact, we examined expression of *sodB* whose expression is RelA-independent (*27*). RNA analysis showed that both RyhB and RyhBm repressed *sodB* expression indicating that mutating GGAG had no effect on the core domain of RyhB (Figure S8A). Yet, RelA mediated repression regulation of *sodA* by RyhBm was null (Figure 6C). As the target mRNA *sodA* harbors an intact RelA binding site, RelA regulation of basepairing RNAs depends on RelA interacting with GGAG carried by sRNAs rather than by mRNA.

In the sRNA/mRNA pair ChiX and *nadE* of *Salmonella*, both *nadE* and ChiX lack the site GGAG (Figure S5). *β*-galactosidase assay of the ChiX/*nadE* pair showed that induction of RelA synthetic activity had no effect on *nadE* repression regulation by ChiX (Figure 6D). In the regulatory pair OxyS and *fhlA* of *E. coli*, only OxyS carries GGAG (Figure S5). *β*-galactosidase assay of the OxyS/*fhlA* showed that upon induction of RelA-synthetic activity, OxyS no longer repressed expression of *fhlA-lacZ* indicating that GGAG site of OxyS is sufficient to enable regulation (Figure 6E). Thus, sRNAs with GGAG trigger RelA function *in vivo*.

## Discussion

### The pathway by which RelA mediates Hfq assembly

In this study we show that Hfq monomer binds RNA in the presence of RelA, challenging the previous concept that Hfq binds RNA only as a hexamer. These results support the notion that RNA binding coincides with Hfq hexamerization which requires initial disassembly of Hfq, and RNA binding to individual Hfq subunits to form a new RNA-bound complex.

Using cross-linking we identified binding of labeled RyhB or *sodA* to one Hfq monomer. The binding was visible only in the presence of RelA, indicating that RelA stabilized the initially unstable complex of RNA associated with Hfq monomers. By mixing limiting amounts of wild type Hfq and high levels of Hfq mutant with RNA that can be bound only by wild type we discovered that preliminary RNA binding to Hfq monomers is necessary and sufficient for RelA to initiate Hfq assembly thereby forming mixed oligomeric sub-unit complexes. Together, these results led us to propose that RelA by stabilizing the originally unstable complex of RNA bound to an Hfq monomer enabled the attachment of additional subunits to form hexamers (see model Figure 7A). The proposed mechanism assumes that RelA is effective in sub-stoichiometric amounts relative to Hfq, which in turn corresponds to the previously reported low intracellular concentration of RelA (*37*).

**Figure 7.**
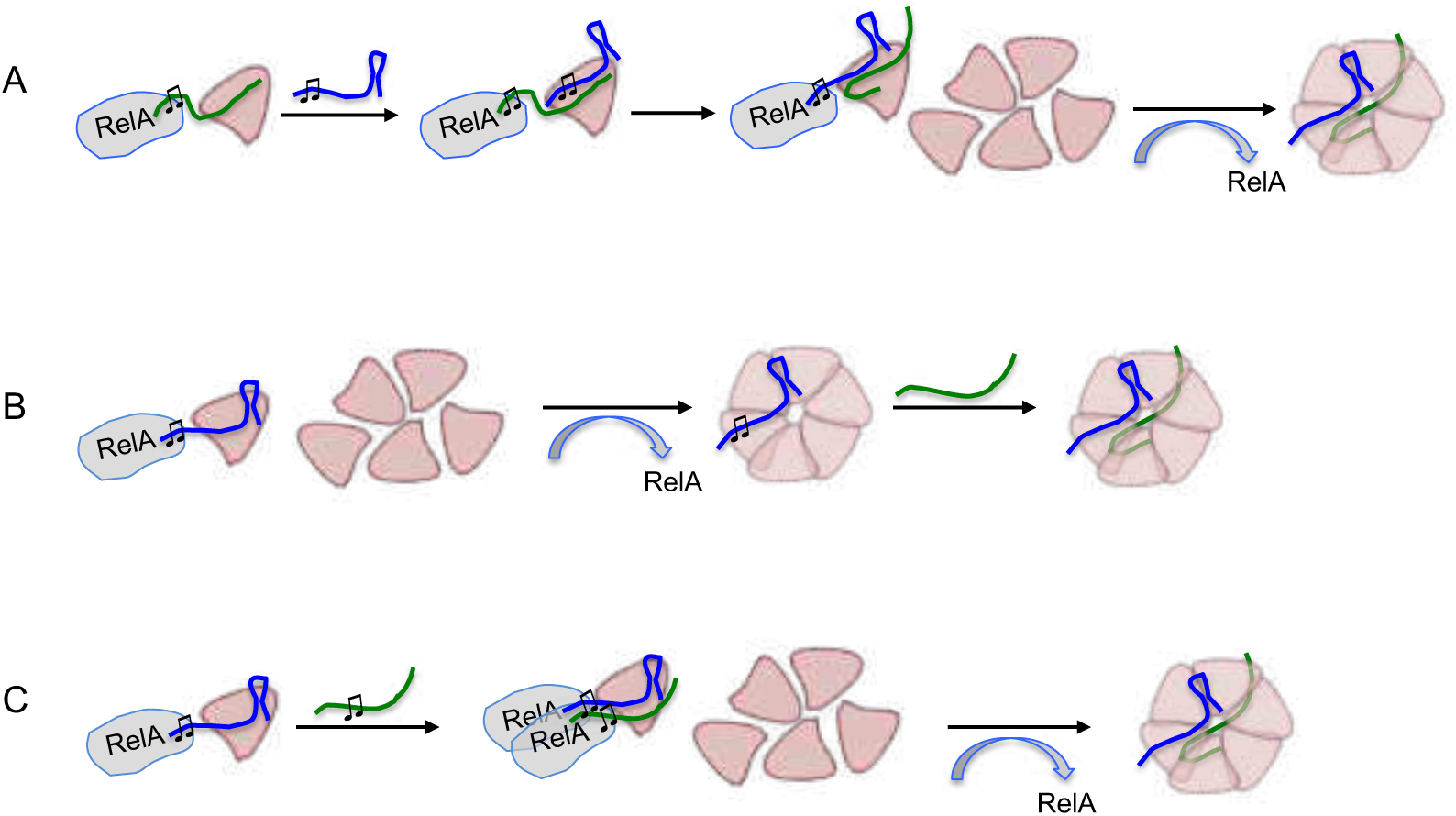
RelA mediated hexamerization of Hfq requires an initial binding of RNA to Hfq monomer. (**A**) RelA binds and stabilizes an unstable complex of mRNA-Hfq monomer leading to subsequent sRNA binding while RelA switches from SD of the mRNA to the GGAG site in the sRNA promoting further stabilization of the complex and enabling Hfq hexamerization. (**B)** RelA stabilizes an unstable complex of sRNA-Hfq monomer by binding the GGAG site in the sRNA leading to Hfq hexamerization followed by target mRNA binding. (**C**) RelA stabilizes an unstable complex of sRNA-Hfq monomer by binding the GGAG site in the sRNA. The binding of SD site in the mRNA possibly by RelA dimer promotes further stabilization and ultimately Hfq hexamerization. sRNA (blue); mRNA (green); Hfq subunits (light brown); music note (♫) symbolizes the GGAG site. In all cases, sRNA mediated mRNA regulation is expedited by RelA binding of sRNAs.

### RelA mediated Hfq assembly requires an initial binding of RNA to Hfq

The use of Hfq mutants demonstrated that RelA mediated Hfq assembly requires an initial binding of RNA to Hfq monomers. In these experiments, Hfq proximal (K56A and D9A) and distal (I30D and G29A) face mutants were incubated with proximal or distal RNAs. RelA failed to induce the RNA binding activity of low levels (5nM) of Hfq proximal mutants presented with proximal face RyhB sRNA (Figure S9A lanes 7, 10) and Hfq distal mutants presented with distal face *sodA* RNA (Figure S9B lanes 13, 16). In contrast, RelA induced the RNA binding activity of low concentrations of Hfq mutants presented with RNAs capable of binding the opposite face of the mutation (Figure S9A lane 13 and S9B lanes 7,10). Interestingly, we noticed that only low concentrations of RelA (25nM) induced the RNA binding activity of Hfq:G29A distal mutant presented with RyhB. Since the RNA binding affinity of G29A is significantly low as compared to Hfq wild type (Figure S9C compare lanes 3, 9, 15) we suspect that high RelA levels (200 nM) competed with Hfq:G29A for the RNA (Figure S9C; lanes 17,18).

Cross linking of Hfq proximal and distal face mutants presented with proximal and distal RNAs showed that while in the absence of RelA, the oligomerization pattern of Hfq mutants presented with *sodA* or RyhB was similar (Figure 10ABCD lanes 1-6), RelA facilitated further oligomerization of Hfq proximal mutants in the presence of distal RNA (Figure S10AB lanes 7-12) and distal mutants presented with proximal RNA (Figure S10CD lanes 7-12), indicating the importance of initial Hfq RNA binding for RelA mediated subsequent oligomerization of Hfq.

### RelA N-terminal domain binds RNA

Deletion and mutational analyses revealed that RelA N-terminal domain is responsible for mediating Hfq-sRNA based target gene regulation. Specifically, two single point mutations in RelA (C289Y and T298I) that clustered in one helix of the RelA N-terminal domain failed to enable repression of RyhB target gene fusions. Unlike wild type RelA that facilitated RNA binding of low concentrations of Hfq by triggering Hfq oligomerization, RelA C289Y and T298I mutants showed no effect on Hfq RNA binding nor they affected Hfq oligomerization. As these RelA mutants produce (p)ppGpp similar to wild type, (p)ppGpp production and RelA mediated basepairing RNA regulation, although both reside in the N-terminal domain were found to be distinct functions. Previously, using a highly sensitive binding assay, we found that incubation of His-tagged RelA-CTD purified from wild type *hfq* cells (a gift from G. Glaser) with RyhB resulted in residual binding of Hfq to RyhB. As RelA-CTD was co-purified with Hfq, we suggested that this portion of the protein might also act as a functional partner of Hfq (*27*). Our current *in vivo* and *in vitro* genetic and biochemical studies demonstrated the importance of the RelA N-terminal domain for Hfq assembly as opposed to its C-terminal domain. Therefore, we suspect that RelA-CTD purified from wild type Hfq cells was contaminated with Hfq due to the presence of 24 histidine residues at its C-termini (*38*). Unlike the intact RelA protein, which was further investigated by its purification from Δ*hfq* cells, we did not explore the function of this domain any further.

### RelA binds RNAs with GGAG

Co-immunoprecipitation using *α*RelA antibody to detect whether Hfq was precipitated in a complex with RelA further confirmed that *in vivo*, RNA links between RelA and Hfq forming a complex. As Hfq did not precipitate with RelA:C289Y mutant, it strongly supported the notion that RelA binds RNA that is bound by Hfq. Further RelA-RNA binding assays demonstrated that RelA binds RyhB with high affinity compared to RelA’s binding affinity for *sodA* and foot printing revealed that RelA binds and therefore protects a specific sequence of GGAG. RNA mutants in which GGAG sequence was changed or deleted rendered RelA inactive *in vitro*; unable to promote Hfq assembly, as well as *in vivo*; incapable of assisting in basepairing regulation by Hfq.

Quantitative RT-PCR analysis of the RNA bound to RelA during Co-IP revealed that sRNAs carrying GGAG sequence including RyhB, SraC and McaS were bound by RelA but not by RelA:C289Y (Figure S11). The calculated copy number of these sRNAs was 6×10^4^, 102 and 309-fold more in lysates of wild type *relA* than in lysate of *relA*:C289Y. The copy number of MgrR and MicC sRNAs that lack the GGAG sequence was identical in both *relA* and *relA*:C289Y. Similarly, the copy number of *sdhC* and *sodA* mRNAs although carry GGAG was also identical in *relA* and in *relA*:C289Y, further confirming that RelA binds a specific class of sRNAs.

### RelA binding of sRNAs vs. mRNAs

Our *in vitro* experiments show that the affinity of RelA to sRNA is higher than its affinity for mRNA carrying the same GGAG sequence. Our *in vivo lacZ* assays demonstrate that RelA binding to sRNAs is essential for target gene regulation. The preference of RelA observed *in vitro* for sRNAs with GGAG to mRNAs with the same sequence is not clear. As the GGAG sequence resembles the Shine-Dalgarno (SD) sequence present in many mRNA targets, it is possible that *in vivo*, RelA binds the SD sequence of mRNAs. In this scenario, RelA binds and stabilizes an unstable complex of mRNA-Hfq monomer leading to subsequent sRNA binding while RelA switches from SD of the mRNA to the GGAG site in the sRNA promoting further stabilization of the complex and enabling Hfq hexamerization. Alternatively, RelA stabilizes an unstable complex of sRNA-Hfq monomer by binding the GGAG site in the sRNA leading to Hfq hexamerization followed by target mRNA binding, or the complex of RelA-sRNA-Hfq is further stabilized via binding of the target mRNA SD site by a second RelA leading to Hfq hexamerization. In either case, sRNA mediated mRNA regulation is expedited by RelA binding of sRNAs (see model Figure 7).

Moreover, comparing RyhB and RyhBm levels in *relA*^+^ and *relA*^-^ strains indicate that sRNA binding by RelA results in sRNA stabilization (Figure S8B). The level of the wild type RyhB RNA was higher in *relA*^+^ than in *relA*^-^, whereas the levels of RyhBm in which RelA binding site was changed were similar in both *relA*^+^ and *relA*^-^. RyhB is unstable in an *hfq*^-^ strain as Hfq protects it from degradation by RNase E (*27, 39*). As the levels of RyhB increase in *relA*^+^, RelA stabilization of the unstable complex of RNA and Hfq is due to sRNA stabilization by Hfq or both RelA and Hfq.

### RelA GGAG sRNA binding and (p)ppGpp productions are mutually exclusive functions

An interaction between RelA-like protein and RNA was documented previously for RelQ (*40*). The authors showed that the small alarmone synthetase RelQ from the Gram-positive pathogen *Enterococcus faecalis* bound mRNAs at AGGAGG sites. The enzymatic activity of *E. faecalis* RelQ was inhibited by mRNA binding, and addition of (p)ppGpp counteracted the inhibition. Because (p)ppGpp synthesis and (p)ppGpp binding were mutually incompatible with RelQ:RNA complex formation, it was proposed that RelQ enzymatic and RNA binding activities are subject to allosteric regulation.

Our data indicate that the N-terminal domain of RelA contains two distinct functions; (p)ppGpp synthesis and RNA binding. *In vitro* and *in vivo* (p)ppGpp production by wild type RelA decreased when presented with RNA carrying GGAG. In contrast the presence of sRNAs lacking this sequence had no effect on RelA (p)ppGpp synthetase activity. The synthetase activity of RelA mutant (C289Y) that is unable to bind RNA remained unaffected, indifferent to any kind of RNA. Similar to RelQ, we suspect that these two functions that reside within the same N-terminal domain although in different positions are mutually exclusive because of allosteric inhibition.

*β*-galactosidase assays of target genes carried out with and without serine hydroxamate to induce (p)ppGpp production showed that repression by sRNAs carrying GGAG (i.e. OxyS and RyhB) was impaired upon induction of RelA (p)ppGpp synthetic activity. In contrast, repression of *nadE-lacZ* by ChiX of which both lack the sequence GGAG was unaffected by induction of RelA synthetic activity. Finally, changing the GGAG site in RyhB (RyhBm) demonstrated that RelA triggers Hfq basepairing regulation by interacting with sRNAs.

### The physiological conditions leading to RelA regulation of basepairing RNAs

*In vivo* estimation of Hfq concentration showed that *relA*^+^ and *relA*^-^ strains harbor approximately 8 µM and 6.5 µM Hfq hexamers, respectively (*41*). However, the absolute concentration of Hfq is not indicative of Hfq availability. Co-IP studies have revealed thousands of Hfq-bound RNAs and overexpression of Hfq-dependent sRNAs resulted in the sequestration of Hfq and thus in Hfq depletion (*13, 42–44*). Here we show that RelA enables binding of RNAs by otherwise ineffective amounts of Hfq *in vitro*, and facilitates Hfq mediated basepairing regulation of specific sRNA/mRNA pairs *in vivo*, indicating that under specific conditions and/or environments, Hfq availability is inadequate.

Our data indicate that RelA is specific to sRNAs with GGAG and that it affects not all sRNA/mRNA pairs. However, what distinguishes the groups is unclear. Conceivably, the affinity of Hfq for the RelA-independent class of sRNAs and/or mRNAs is high and therefore also low levels of Hfq are effective. For example, the binding affinity of the GGAG sequence lacking sRNAs such as ChiX, DsrA, RprA, MgrR, MicA, MicC, MicF and SpoT42 was estimated by filter binding or gel mobility shift assays to be 0.21 nM, 0.54 to 23 nM, 25 nM, 0.48 nM, 2.3 nM, 3.3 nM, 1.7 nM and 20 nM, respectively (*17, 24, 45-47*). Also, ChiX, MgrR and DsrA that lack the GGAG sequence were reported to be better competitors for binding Hfq than RyhB, McaS or CyaR sRNAs (*46*). Interestingly, both MgrR and ChiX carry in addition to poly(U) tail three and four ARN motifs, respectively (*25, 45*). Deleting these motifs led to a significant loss of stability of both sRNAs, while adding these motifs to RyhB increased its stability (*25*). Thus, indicating that additional binding of the sRNAs to Hfq distal site by the ARN motifs apart from the proximal site interaction results in enhanced affinity and hence stability of the sRNAs.

In investigating the occurrence of sRNAs with GGAG, we identified 26 sRNAs with this sequence from a group of 86 (Figure S12). Using a combination of the Clustal omega (https://www.ebi.ac.uk/Tools/msa/clustalo/) and Genedoc software we identified 19 sRNAs in which the GGAG site was conserved in several bacterial species (Figure S13). Of the 26 sRNAs, many are expressed during stationary phase and/or in minimal media such as McaS, RybB, RydB, RyfD, RyeA/SraC, RyhB and GadY. A few are generated from the 3’UTR of protein coding genes (*glnA*, *kilR*, *malG*, *allR*). Intriguingly, a significant number of the sRNAs belongs to type I toxin-antitoxin (TA) systems (RalA, SibA, SibC, SibE, SokC and SokE). Among these TA systems, only RalA is known to be stabilized by Hfq (*48*). It may be that the Hfq binding affinity of these sRNAs is extremely low and almost undetectable, thus requiring RelA assistance. Alternatively, RelA binding of these sRNAs plays a regulatory role in the absence of Hfq. As RelA synthetase activity is incompatible with RelA regulation of base-paring RNAs, it is intriguing to speculate that upon normal growth conditions, RelA facilitates repression of the TA systems to decrease toxicity, whereas under conditions of amino acid starvation, RelA indirectly leads to an increase in expression of the toxin genes of the TA systems thereby modulating primary metabolic pathways.

## Methods

### Bacterial strains and plasmids

Strains, plasmids and primers used in this study are listed in Tables S1, S2 and S3. Bacteria were grown routinely at 37°C in Luria-Bertani (LB) medium. Ampicillin (amp, 100µg/ml), kanamycin (kan, 40µg/ml), chloramphenicol (cm, 30µg/ml) and tetracycline (tet, 10µg/ml) were added where appropriate.

### Strain construction

Chromosomal gene deletion mutants were carried out using the kanamycin and chloramphenicol cassettes of pKD4 and pKD3, respectively (*50*) as described previously (*51*). The chromosomal deletions were transferred into fresh genetic background by transduction using the P1 bacteriophage. To construct *ΔrelA::kan*, primers 2187 and 2188 were used to replace 4kb region encompassing *relA1* with 1.2 kb of kanamycin cassette. *ΔryhB::cam* was constructed using primers 618 and 619. *Δhfq::cam* was constructed using primers 2383 and 2384. To generate *ΔrelA::frt, Δhfq::frt* and *ΔryhB::frt*, the antibiotic resistance genes were removed using pCP20 (*50*).

### Plasmid constructions

The *relA* gene was amplified from *E. coli* MC4100 chromosomal DNA using the primers 2530 and 2345. The PCR product was digested with the *Pst*I and *Hind*III restriction enzymes and ligated downstream of P_BAD_ of p15A vector generating pRelA. To construct pRelA-ΔCTD (2971-2972) and pRelA-ΔNTD (2973-2974), whole plasmid PCR was carried out using primers as above and pRelA as template. Point mutations in the *relA* gene including C289A (3039-3040) and Q264E (3134-3135) were introduced in pRelA plasmids using the Gibson cloning method (*52*). The pRelA:C289Y;Y290C plasmid was generated by introducing the Y290C mutation (3036-3037) in the pRelA:C289Y plasmid (obtained by random mutagenesis). For purification of the RelA protein, we cloned the *relA* gene in pET-15b vector by the Gibson cloning method. To this end primers 3031-3032 were used to amplify pET-15b vector and primers 2999-3000 were used to amplify the *relA* gene. RelA point mutations; C289Y (3069-3070), C289Y;Y290C (3036-3037) and C289A (3039-3040) were cloned in pET-15b using the same method (Gibson). For Hfq protein purification, the *hfq* gene was amplified using the primer pairs 2687-2351 and cloned in the pET-15b vector using *Nco*I/*BamH*I restriction enzymes (PT7-Hfq). Hfq point mutations; D9A (3168-3169), K56A (3048-3049), G29A (3170-3171) and I30D (3054-3055) were introduced in PT7-Hfq using Gibson cloning method. P_L_-RyhBm was constructed by whole plasmid PCR using primers 3266-3267 and P_L_-RyhB as template. The *chiX* gene fragment of SL1344 was amplified using primers 2594 and 2595 and subcloned downstream of P_BAD_ into the unique *Eco*RI and *Hind*III sites of pJO244. To construct *nadE*-*lacZ* translational fusion in pSC101 (pBOG552), *nadE* 5’-end fragment carrying 166 nucleotides from –327 upstream of the AUG initiation codon to +164 was amplified using primers 2907-2908 and subcloned into the unique *Eco*RI and *Bam*HI sites of pBOG552. To construct *sodA-lacZ* translational fusion, the *sodA* 5’-end fragment carrying 391 nucleotides from -293 upstream of the AUG initiation codon to +98 was amplified from MC4100 chromosomal DNA by PCR using oligonucleotides 2927 and 2928 and subcloned into the unique *Eco*RI and *Bam*HI sites of pRS552 (*53*). To construct MC4100 *sdhC-lacZ* translational fusion, the *sdhC* 5’ end fragment carrying 319 nucleotides from -281 upstream of the AUG initiation codon to +38 was amplified from MC4100 chromosomal DNA by PCR using oligonucleotides 2491 and 2505 and subcloned into the unique *Eco*RI and *Bam*HI sites of pRS552. The *lacZ* fusions were then recombined onto λRS552 and integrated into the attachment site of *relA*^+^*ΔryhB*::*frt* (A-506)*, ΔrelA::frt* (A-1036) and *ΔrelA::frt ΔryhB::frt* (A-1046).

### Random mutagenesis

Random mutagenesis of the *relA* gene (pRelA; p15A) was carried out using hydroxylamine as described before (*54*). The mutagenized plasmids were transformed into MC4100 Δ*relA* strains carrying *sdhC-lacZ* chromosomal fusion and a plasmid expressing RyhB (P_L_-RyhB; ColE1). Blue versus white colonies were selected on LB plates containing 40µg/ml of 5-bromo-4-chloro-3-indolyl-β-D-galactopyranoside (X-Gal) and 0.1% arabinose.

### Scarless point mutations in the chromosome

Chromosomal scarless point mutations within *relA* were carried out as described in (*55*). Briefly, the *tetA-sacB* cassette from the XTL634 strain chromosome was amplified using the primer pairs 3112-3113 carrying sequences homologous to the *relA* gene. The PCR product was inserted into *relA*^+^*hfq*^+^ (A-397) and *relA*^+^Δ*hfq*::*frt* (A-950) generating the *relA*:*:tetA-sacB* strain. Next, PCR product generated using primer pairs 3114-3115 that amplify the C289Y mutation from the (p_BAD_-RelA; p15A) was used to transform *relA*:*:tetA-sacB*. Colonies sensitive to tetracycline were selected on fusaric acid containing plates.

### β-galactosidase assays

Strains as indicated were grown for 16-18h in M9 minimal medium containing 0.4% glycerol, 0.04% glucose and 0.2% arabinose for induction of RelA expression. The cultures reached OD600 of ∼ 0.3-0.4. LacZ assays were carried out as previously described (*54*). To determine the effect of RelA mediated regulation of RyhB, ChiX and OxyS target genes under amino acid starvation, the strains as indicated were grown for 16-18 h in MOPS minimal medium supplemented with 0.4% glycerol and 0.04% glucose and 0.2% arabinose (for ChiX induction) in either the presence or absence of 500µg/ml of Serine hydroxamate (SHMT). LacZ assays were carried out as above.

### RelA protein purification

BL21-DE3 Δ*relA* Δ*hfq* strains carrying pET-15b plasmids expressing the 6X His tagged RelA wild type, RelA:C289Y, RelA:C289Y;Y290C and RelA:C289A grown in 1lt of LB at 37°C to OD600 of 0.5-0.6 were treated with 1mM IPTG and continued to grow to OD600 of 2. The pellets were washed once in 1X PBS and then dissolved in 30ml of cold buffer A {10mM imidazole, 500mM NaCl, 200mM NaH_2_PO_4_ (pH 7.4)} along with 50µl each of PMSF, DNase I (0.1mg/ml) and 1M MgCl_2_. Following vortex, homogenization and lysis in micro fluidizer, the supernatant was collected by centrifugation and loaded onto His Trap Chelating HP column (1ml) in the Ni-AKTA Prime machine. The column was washed with washing buffer {400 mM imidazole, 500 mM NaCl, 20 mM NaH_2_PO_4_ (pH 7.4)} and fractions of 8 ml were collected in the fraction collector. Fractions showing the maximum amounts of RelA protein were collected in a snakeskin dialysis bag and subjected to cleavage by TEV protease (overnight, 4° C, dialysis in buffer A). The cleaved protein was passed through the column again and eluted with washing buffer. All Fractions were collected and subjected to dialysis overnight in RelA buffer {50mM Tris-Ac (pH 8.5), 10mM potassium phosphate buffer (pH 8.5), 10mM EDTA, 1mM DTT, 25% glycerol}.

### Hfq protein purification

BL21-DE3 Δ*hfq* carrying pET-15b plasmids expressing Hfq wild type, Hfq K56A, Hfq I30D, Hfq D9A and Hfq G29A grown in 1lt of LB at 37°C to OD600 of 0.5-0.6 were treated with 1mM IPTG and continued to grow to OD600 of 2. The pellets were washed once in 1X PBS and then dissolved in 50ml of lysis buffer {50mM Tris (pH 8), 1.5M NaCl, 250mM MgCl_2_, 1mM β-mercaptoethanol} along with 50µl each of PMSF, DNase I (0.1mg/ml) and 1M MgCl_2_. Following vortex, homogenization and lysis in micro-fluidizer, the supernatant was collected by centrifugation, heated at 85°C for 45 min, clarified by centrifugation and treated with 30µg/ml of RNase A for 1h at 37°C. After RNaseA treatment, the supernatant was loaded onto His Trap Chelating HP column (1ml) in the Ni-AKTA Prime machine. The column was washed with washing buffer {50mM Tris (pH 8), 1.5M NaCl, (pH 7.4), 0.5mM β-mercaptoethanol} and 8 ml fractions were collected in the fraction collector. Fractions showing the maximum amounts of Hfq protein were collected in a snakeskin dialysis bag and subjected to overnight dialysis in Hfq buffer {50mM Tris (pH 7.5), 50mM NH_4_Cl, 1mM EDTA, 10% glycerol}.

### Primer extension

Total RNA (15µg) extracted using Tri reagent (Sigma) from strains as indicated was incubated with end labeled *sodB* specific primer (810) at 70°C for 5 min, followed by 10 min in ice. The reactions were subjected to primer extension at 42°C for 45 min using 1 unit of MMLV-RT (Promega) and 0.5mM of dNTPs. Extension products were analyzed on 6% acrylamide 8M urea-sequencing gels.

### Northern analysis

RNA samples (15µg) isolated from strains as indicated were denatured for 10 min at 70°C in 98% formamide loading buffer, separated on 6% acrylamide 8M urea gels and transferred to Zeta Probe GT membranes (Bio-Rad laboratories) by electroblotting. To detect RyhB, the membrane was hybridized with end labelled RyhB primer (470) in modified CHURCH buffer (1mM EDTA, pH 8.0, 0.5M NaHPO4, pH 7.2, and 5%SDS) for 2 h at 45°C and washed as previously described (*56*).

### *In-vitro* RNA synthesis

DNA templates for RNA synthesis: RyhB, 90 nucleotides used for crosslinking and foot-print were amplified using primers 678-567; RyhB 50 nucleotides used for EMSA was generated using primers 678-3265; RyhB mutant, 90 nucleotides used for crosslinking was generated using primers 3272-567; *sodA* 210 and 56 nucleotides used for EMSA were generated using primers 1764-1765 and 3209-3211 respectively; *sodA* 98 nucleotides used for crosslinking and foot-printing was generated using primers 1764-3203 and *sodA*-ΔSD 47 nucleotides used for crosslinking was generated using primes 1764-3198. The 98 nucleotides *sodAm* template for foot-printing was obtained from TWIST Bioscience. RNAs were synthesized with phage T7 RNA Polymerase (25 units, NEB) in 50µl reaction containing 1X T7-RNA Polymerase buffer, 10mM DTT, 20 units of recombinant RNase inhibitor, 500µM of each NTP, and 300 ng of T7 promoter containing template DNA at 37°C for 2h, followed by 10 min at 70°C. Thereafter, Turbo DNase was added and the reactions were incubated at 37°C for 30 mins. The RNA was purified by phenol-chloroform extraction and then precipitated using 0.3M ammonium acetate, ethanol and quick precip. Labeled α-P^32^ ATP RNAs were generated using low concentrations of unlabeled ATP (20 µM).

### Electrophoretic mobility shift assay

Reactions (10µl) in binding buffer C {50mM HEPES (pH 7.5), 10mM MgCl_2_, 100mM NH_4_Cl and 1.5mM DTT} carrying labeled RNA and/or Hfq and/or RelA as indicated were incubated at 22°C for 10 min and analyzed on 4% native gels using native loading buffer.

### Protein crosslinking assay

Purified proteins along with RNAs as indicated in the figures were incubated in binding buffer C {50mM HEPES (pH 7.5), 10mM MgCl_2_, 100mM NH_4_Cl and 1.5mM DTT} at 22°C for 10 min followed by UV crosslinking for 5 min (254 nm 20000 µJ/ cm^2^). Samples were collected before and after the addition of 0.2% glutaraldehyde at the time points indicated in the figures. Reactions were stopped by adding 200 mM of fresh glycine followed by heating at 95°C for 10 min in sample loading buffer. The proteins were separated in 4-20% MOPS gradient gel. Hfq was detected by Hfq specific antibody (western).

### UV crosslinking assay with RelA

To determine the binding of RNA to RelA, purified RelA (wild type or C289Y mutant) proteins were incubated with 1nM of labelled RyhB or *sodA* at 22°C for 10 min, in binding buffer C followed by UV crosslinking for 5 min as above. Where indicated, 100nM of unlabeled RNA as competitor RNA or RNAse A (100µg/ml for 1h at 37°C) to remove unbound RNA were added. Proteins heated at 95°C for 10 min in sample loading buffer were analyzed in 15% SDS-PAGE.

### DMS and RNaseT1 foot-printing

DMS and RNase T1 foot-printing reactions were carried out as described previously with slight modifications (*41*). Briefly, 0.5pmol of RNA was incubated with 5pmol of RelA (wild type or C289Y mutant) at 22°C for 10 mins, followed by incubation with 0.1, 0.2 or 0.4U of RNase T1 (37°C for 5 min) or 0.3% DMS (25°C for 5 min) in their respective buffers. Reactions were stopped by phenol/chloroform extraction in the presence of 5µg of yeast t-RNA for RNaseT1 and precipitated using 0.5M NaCl and quick precip. Primer extensions to detect the products were carried out using 5’-end labelled RyhB (3273) and *sodA* (3203) primers. The products were separated in 6% acrylamide 8M urea gels.

### Protein Co-immunoprecipitation (Co-IP) assay

Strains as indicated in the figure: *E. coli relA*^+^*hfq*^+^ (A-397), *relA*^+^Δ*hfq*::*frt* (A-950), *relA::C289Y hfq*^+^ (D-1182) and *relA::C289Y* Δ*hfq*::*frt* (D-1172) were grown overnight in M9 minimal medium containing 0.04% glucose and 0.4% glycerol. The pellets (50 OD total) were re-suspended in 1ml of lysis buffer (20mM Tris pH8.0, 150mM KCl, 1mM MgCl_2_, 1mM DTT). Centrifuged at 11200g for 5 mins in 4°C. Thereafter the pellets were flash frozen in liquid nitrogen and thawed on ice. The cells were lysed using 800µl of lysis buffer and 800µl of glass beads (0.1mm) in a bead beater with 30 secs burst and intermittent chilling on ice for a total of 5 min. The lysates were collected by centrifugation while one half of the lysate was treated with 100µg/ml of RNaseA for 1h at 37°C. Immunoprecipitation was carried out with 35µl of rabbit anti-RelA antibody, incubated shaking for 1h at 4°C. Then, 75µl of Protein A Sepharose beads (pre-washed in lysis buffer) were added and the mixture was further incubated with shaking for 1h at 4°C. The beads were washed 5 times with lysis buffer, soaked in 1X SDS sample loading buffer, boiled for 5 mins at 95°C. The samples were analyzed in 4-20% MOPS gradient gel and the proteins detected using rabbit anti-Hfq.

### Isolation of RNA precipitated during Co-IP assay

Co-IP was carried out as above except that the cells were resuspended in lysis buffer containing 20units of RNase inhibitor. The RNA was isolated from the sepharose beads by the TRI-reagent, followed by precipitation with isopropanol and Glycoblue. RNA pellets were further washed with ethanol, air dried and dissolved in 15µl of DEPC.

### Quantitative Real-time PCR (qRT-PCR)

RNA concentrations (obtained after Co-IP extraction) were checked by NanoDrop machine (NanoDrop Technologies). DNA contaminations in the RNA samples were removed by DNase treatment according to the instructions provided by the manufacturer (RQ1 RNase free DNase, Promega). cDNA was synthesized from 2µg of DNA free RNA using MMLV reverse transcriptase and random primers (Promega). Quantification of the cDNA was carried out in the Rotor gene 3000A machine (Corbett) using the real-time PCR SYBR-green mix (Absolute SYBR GREEN ROX MIX, ABgene). Reactions and machine handling were carried out according to manufacturer’s instructions. Genes tested for real time PCR were RyhB (566-3273), MgrR (3305-3306), MicC (3307-3308), SraC (3309-3310), McaS (3313-3314), *sdhC* (3302-3303) and *sodA* (3301-1765). Primer designing was carried out according to the guidelines provided by the IDT PrimerQuest software (https://eu.idtdna.com/PrimerQuest/Home/Index?Display=SequenceEntry). Secondary structure formation within each primer was determined by the IDT OligoAnalyzer software (http://eu.idtdna.com/analyzer/Applications/OligoAnalyzer/). A standard curve was obtained by carrying out PCR with serially diluted *E. coli* MC4100 genomic DNA. Copy number calculation of each gene present in each sample was analyzed by the Rotor-gene analysis software 6.0.

### Western Blotting

To detect Hfq or RelA, protein samples were separated by either 15% acrylamide gel (RelA) or 4-20% MOPS gradient gel (Hfq). Thereafter, the proteins were transferred to nitrocellulose membrane (Genscript) by the Genscript eBLOT L1 fast wet protein transfer system, as suggested by the manufacturer. After transfer, the membrane was incubated at room temperature for 1h (shaking) in blocking solution containing 4% BSA, 4% skim milk and 1X TBST. The membrane was rinsed once with 1X TBST followed by incubation at room temperature for 1h (shaking) in 10 ml of 1X TBST containing 3% BSA and 20µl of rabbit anti-Hfq or anti-RelA antibody. Following three times wash (10 min each) with 1X TBST, the membrane was incubated in HRP conjugated secondary antibody solution (1µl in 10ml of 1X TBST) for 1h with shaking. The membrane was then rinsed once with 1X TBST and the protein were detected by incubation in ECL solution (Advansta Western Bright) for 1 min followed by the use of Image Quant LAS 4000 mini software.

### *In-vitro* (p)ppGpp assay

1µM of purified RelA protein (wild type or C289Y) pre-incubated with or without RNAs in binding buffer C at 22°C for 10 min were further incubated at 30°C for 1h in 1X synthesis buffer {2.5mM GTP, 20mM ATP, 200mM Tris-HCl (pH7.4), 5mM DTT, 50mM MgCl_2_, 50mM NH_4_Cl, 50mM KCl and 10 µCi α-P^32^ GTP}. The products were analyzed by thin-layer chromatography (TLC) with 1.5M of KH_2_PO_4_ (pH 3.4) running buffer. The intensity of GTP and (p)ppGpp spots were measured by the Image Quant software and % of (p)ppGpp production was calculated.

### *In-vivo* (p)ppGpp assay

The intracellular (p)ppGpp accumulation was determined according to the protocol described by (*57*). Briefly, strains grown in LB to OD600 of 0.3 were washed in low phosphate (0.2mM of K_2_HPO_4_) MOPS minimal medium supplemented with all amino acids except serine and inoculated (1:100) in the same medium and grown to OD600 of 0.2. The cells were labelled with 100µCi/ml of {^32^Pi}H_3_PO_4_, by incubation at 37°C for 10 mins, followed by 1h induction of amino acid starvation by the addition of 500µg/ml of Serine Hydroxamate (SHMT). Thereafter, the cells were pelleted, washed with 10mM Tris-HCl (pH 8.0) and resuspended in the same buffer (20µl) followed by lysis using an equal volume of pre-chilled 13M of formic acid with intermittent tapping for 15 min on ice. Cell debris were eliminated by spinning at 13000 rpm for 6 min at 4°C. The collected supernatants were spotted on TLC plates with 1.5M of KH_2_PO_4_ (pH 3.4) running buffer. The intensity of GTP and (p)ppGpp spots were measured by the Image Quant software and % of (p)ppGpp production was calculated.

## Acknowledgments

We are grateful to Sarah Woodson for her advice throughout the study. This work was supported by: the Israel Science Foundation founded by The Israel Academy of Sciences and Humanities (138/18), and the Deutsch-lsraelische Projektkooperation (AM 441/1-1 SO 568/1-1).

## Author Contributions

P.B. performed the experiments and helped wring the manuscript, F.H, M.K. and R.W. helped purifying the proteins, M.E-W. helped in conceptualization of the experiments and reviewing of data. S.A. was responsible for conceptualization of the experiments, reviewing of data, writing the manuscript and funding.

**Figure S1.**
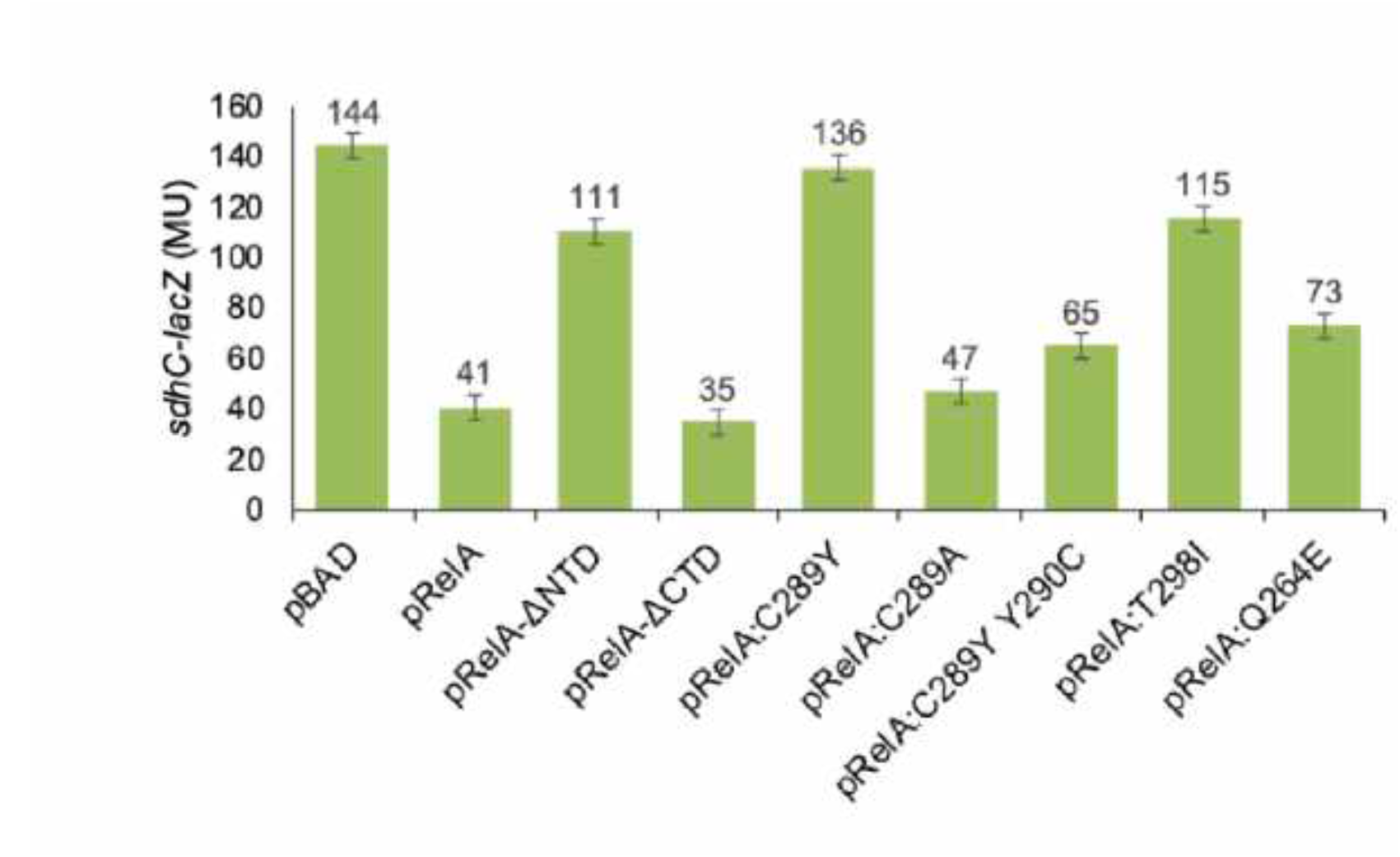
β-galactosidase assay to determine the effect of plasmids encoded RelA alleles on repression of *sdhC-lacZ* target gene fusion by RyhB. Expression of RelA from BAD promoter was induced with (0.2%) arabinose.

**Figure S2.**
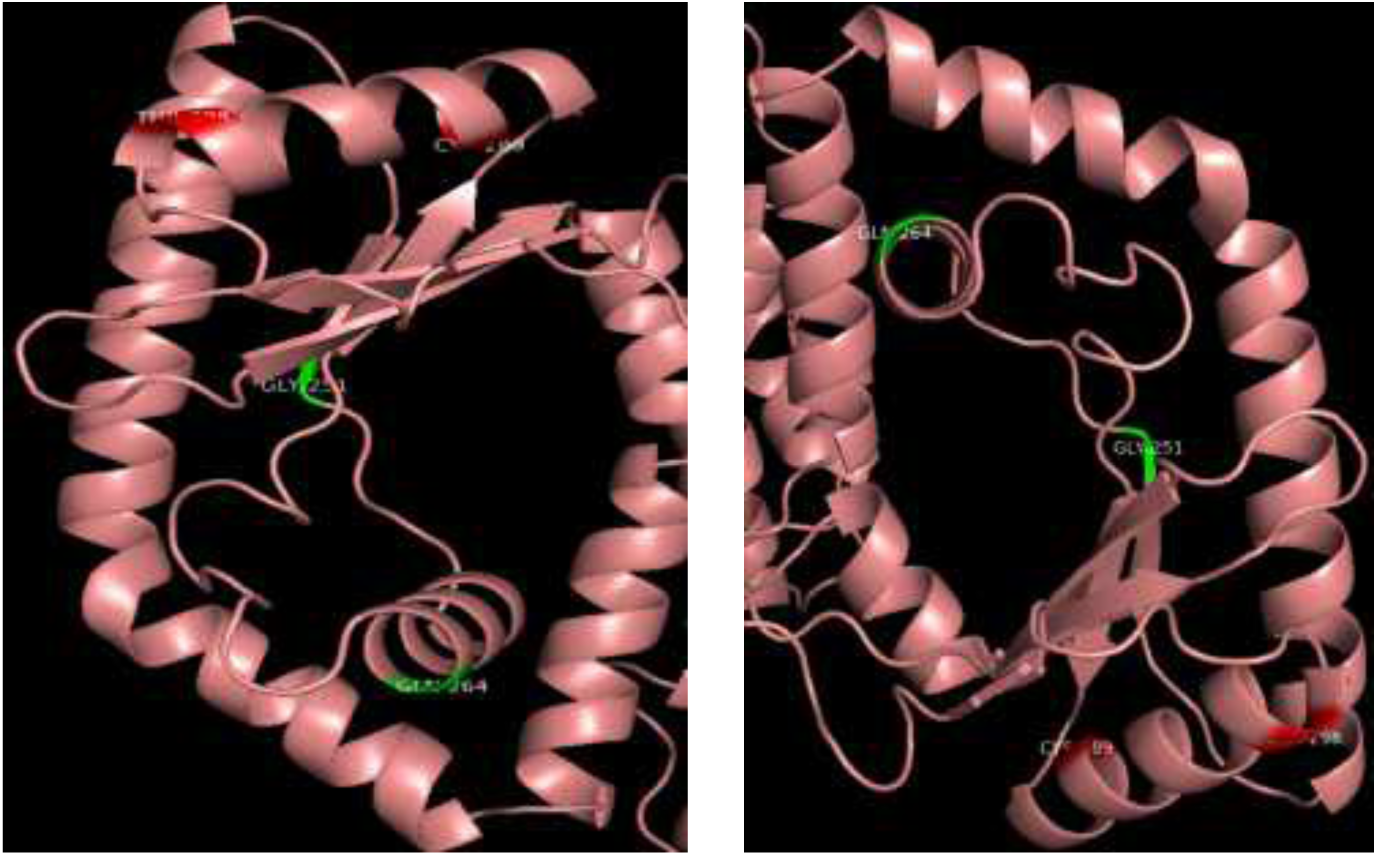
Pymol representation of the amino terminal portion of RelA. The amino acids marked in red (C289 and T298) are essential for RNA binding activity of RelA, while the amino acids marked in green (G251 and Q264) are essential for the (p)ppGpp synthetic activity of RelA. Presented are two forms of display.

**Figure S3.**
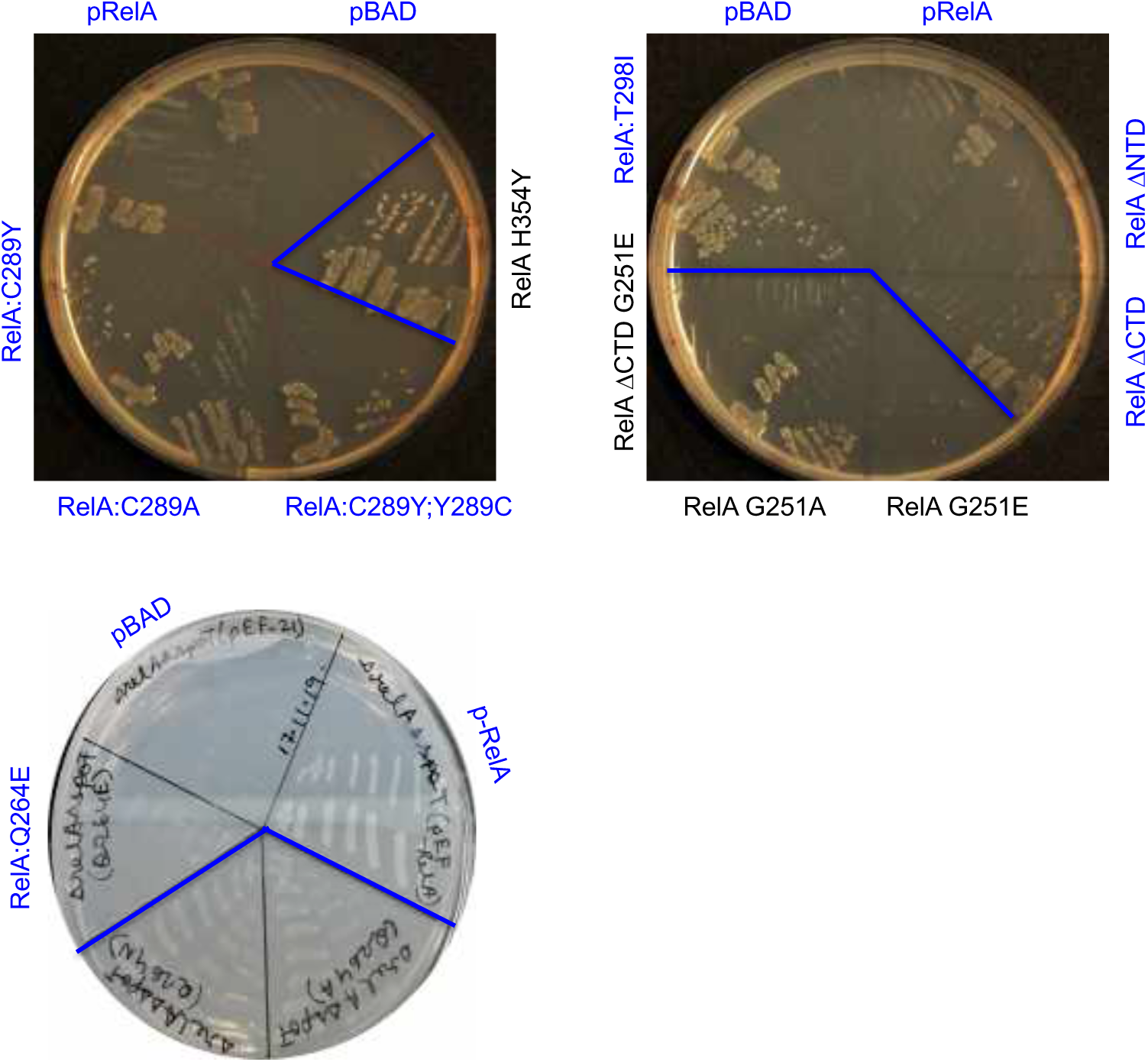
Plasmids expressing RelA enable growth of *E. coli* Δ*relA* Δ*spoT* in M9 minimal medium supplemented with 0.04% glucose, 0.4% glycerol and 0.1% arabinose to induce expression from BAD promoter. The plasmids denoted in blue are relevant for this study.

**Figure S4.**
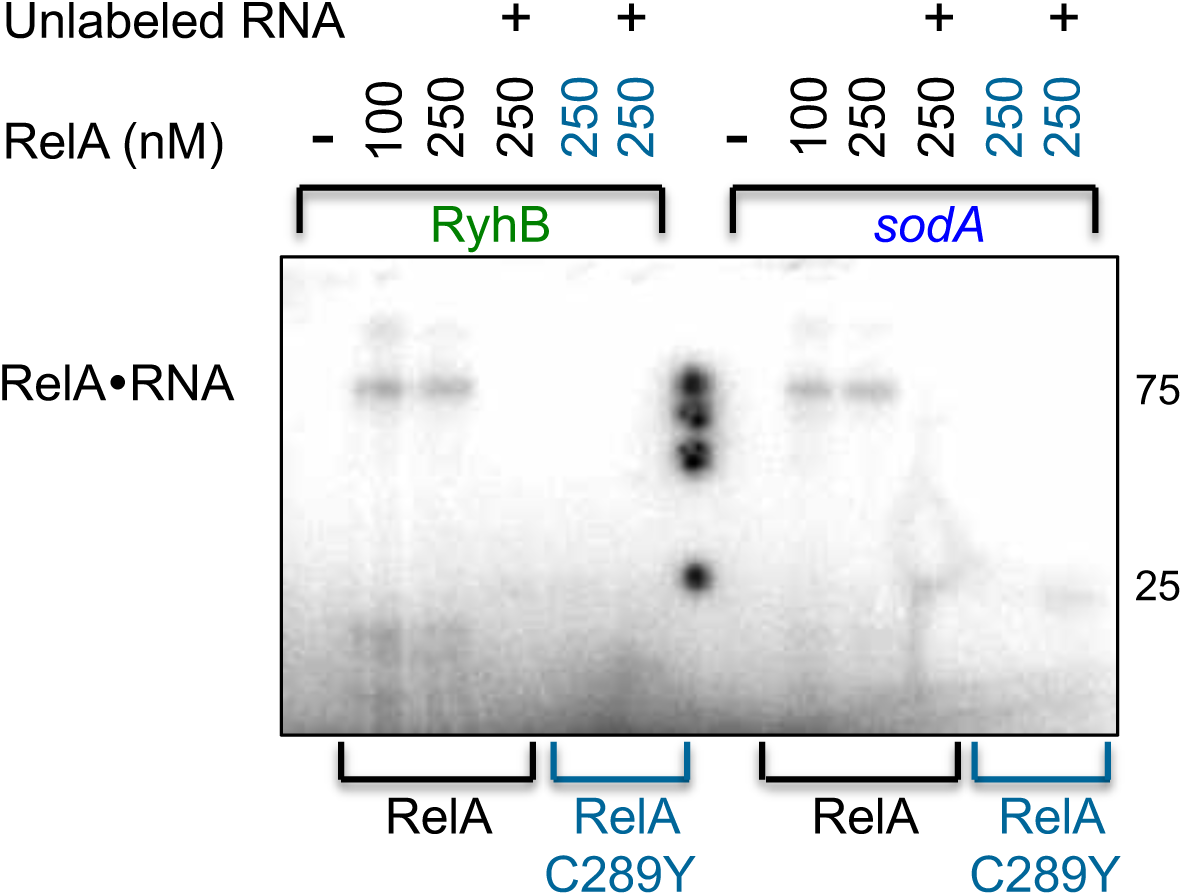
Binding of RyhB and *sodA* by RelA. Wild type RelA or RelA:C289Y incubated with labeled RNAs (1 nM) were UV cross-linked. Then, unprotected RNA residues were digested with 100 µg/ml of RNase A. Competitor unlabeled RNA (100 nM) was added to the reaction mixtures as indicated. The binding products were analyzed by 15% SDS-PAGE. The estimated MW of the RNA•RelA complex is ∼ 85kDa.

**Figure S5.**
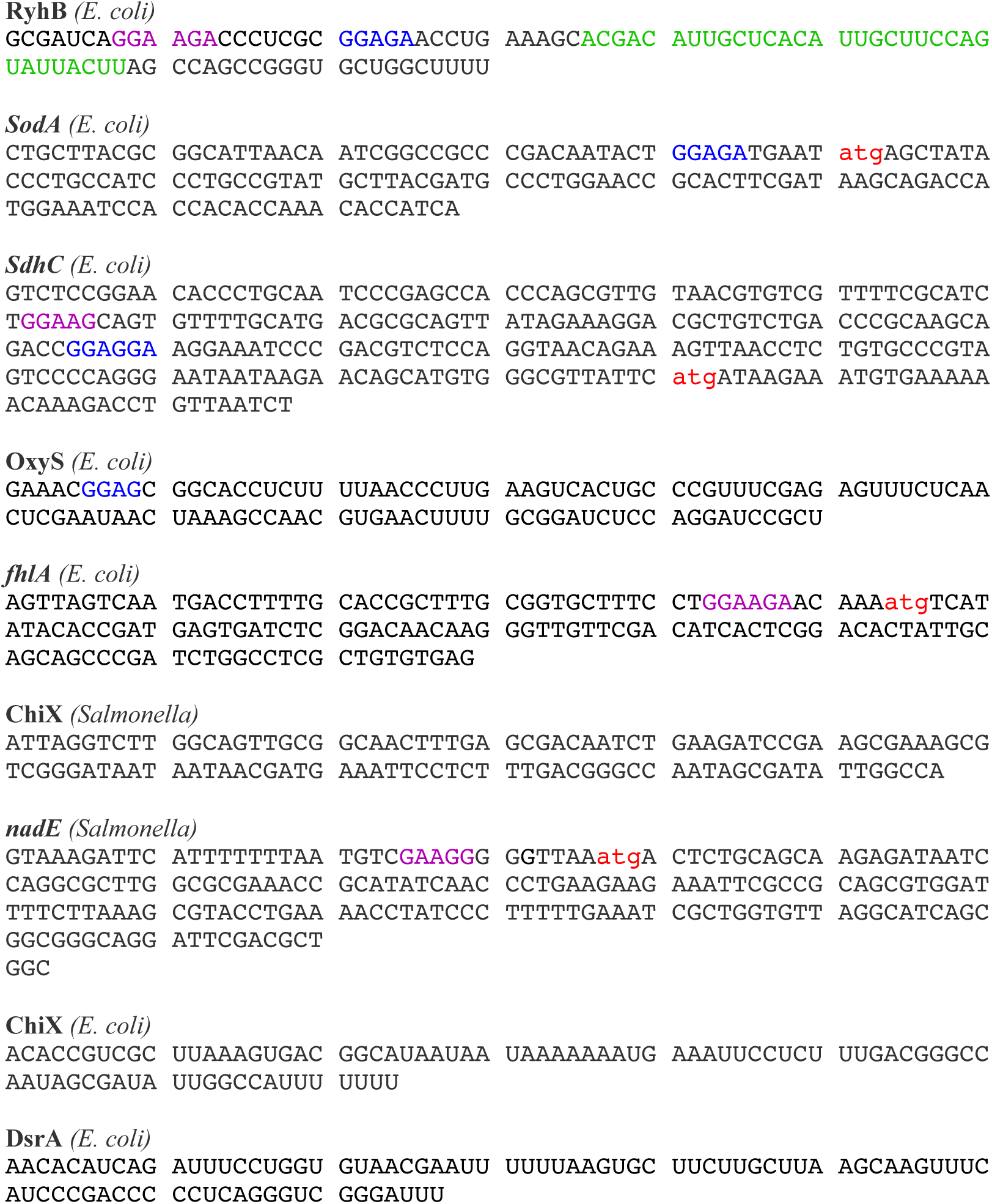
sRNAs and target mRNAs carrying GGAGA. Indicated are AUG (red), GGAGA (blue) and the variant sequences (purple).

**Figure S6.**
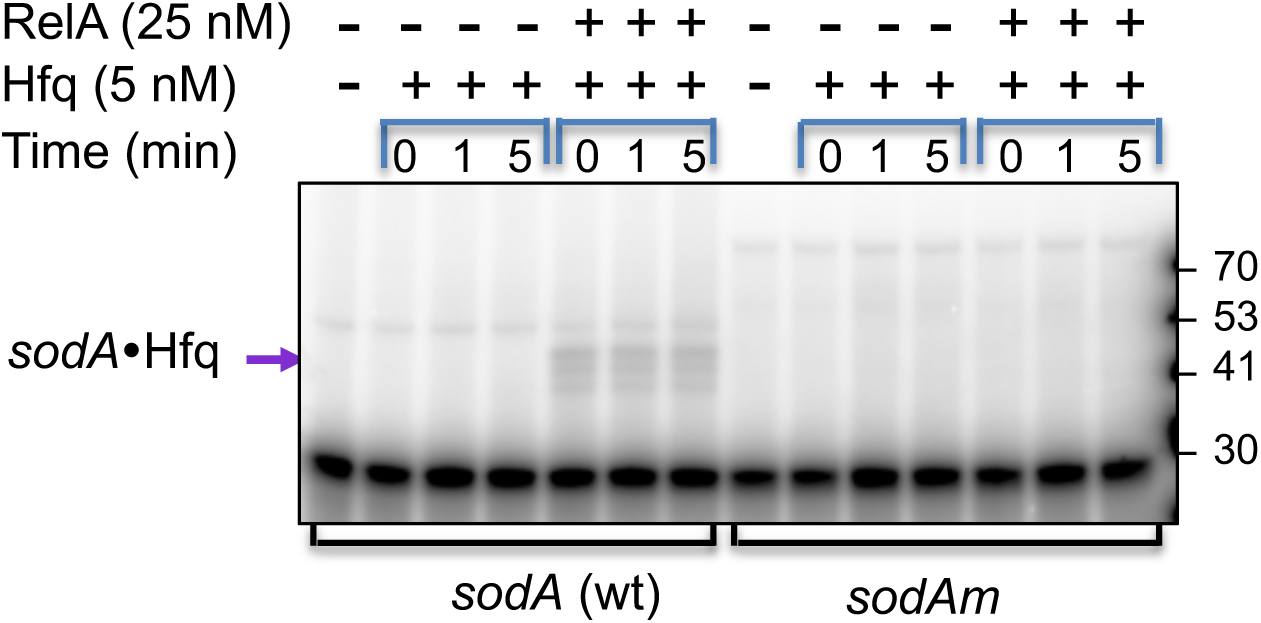
Hfq binding. Reaction mixtures of Hfq incubated for 10 min at 22°C with labeled *sodA* or *sodAm* (1 nM) without or with RelA were UV cross-linked followed by protein crosslinking with 0.2% glutaraldehyde. The cross linking was stopped with 200 mM of fresh glycine and the products analyzed in 4-20% MOPS gradient gel. RNA bound to one Hfq monomer is indicated by the purple arrow on the left side. Note that the addition of RNA alone to low levels of Hfq (5 nM) does not result in Hfq-RNA stable binding. Also, RelA does not induce binding of RNA lacking GGAG to Hfq.

**Figure S7.**
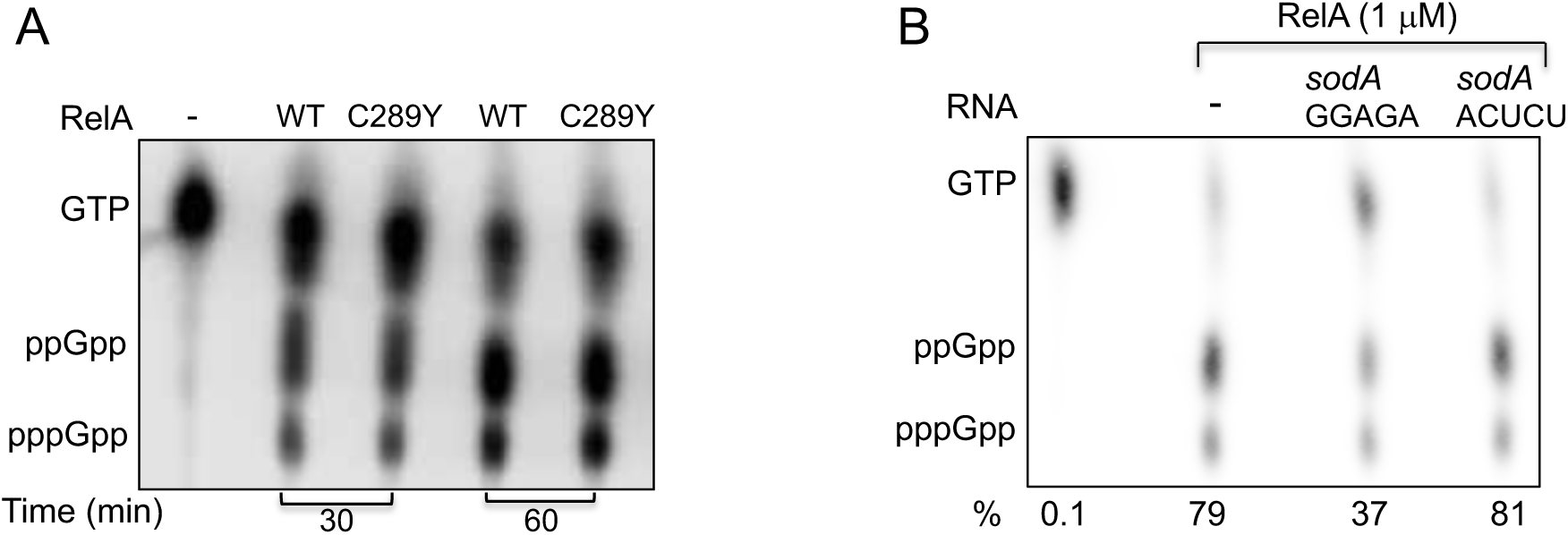
*In-vitro* (p)ppGpp production assays (A) Purified RelA:C289Y produces (p)ppGpp similar to wild type. Purified RelA wild type (WT) and RelA:C289Y mutant (C289Y) were incubated for the times indicated and ppGpp levels were assayed as describe in Material and Methods B. *sodA* carrying an intact GGAGA site inhibits *in-vitro* (p)ppGpp production by RelA. The intensity of the spots was determined by the Image Quant software and percentage of (p)ppGpp production of the total was calculated (% conversion).

**Figure S8.**
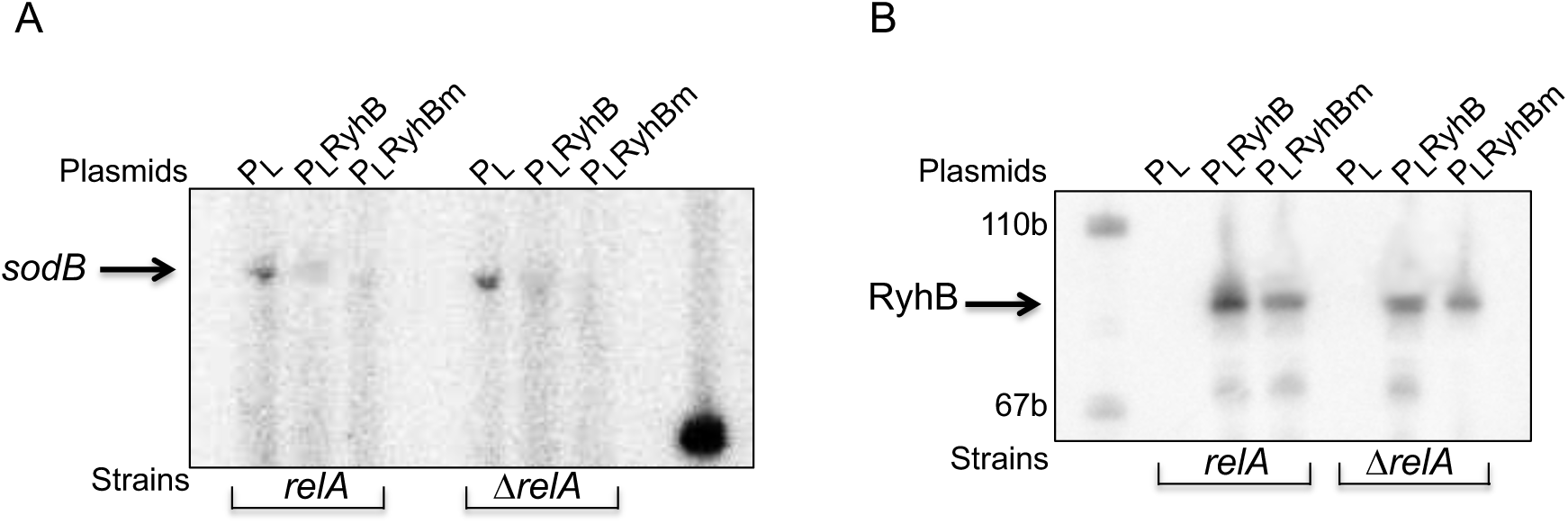
Characteristics of RyhB mutant lacking GGAG (RyhBm) (**A**) Both wild type and RyhB mutant repress *sodB*, a RelA-independent Target. Primer extension using an *sodB* specific primer carried out with RNA isolated from Δ*ryhB,relA*^+^ and Δ*ryhB,*Δ*relA* strains carrying RyhB expressing plasmids (**B**) RelA stabilization of wild type RyhB. RNA as in A was analyzed by northern to detect the levels of wild type and RyhB mutant (RyhBm).

**Figure S9.**
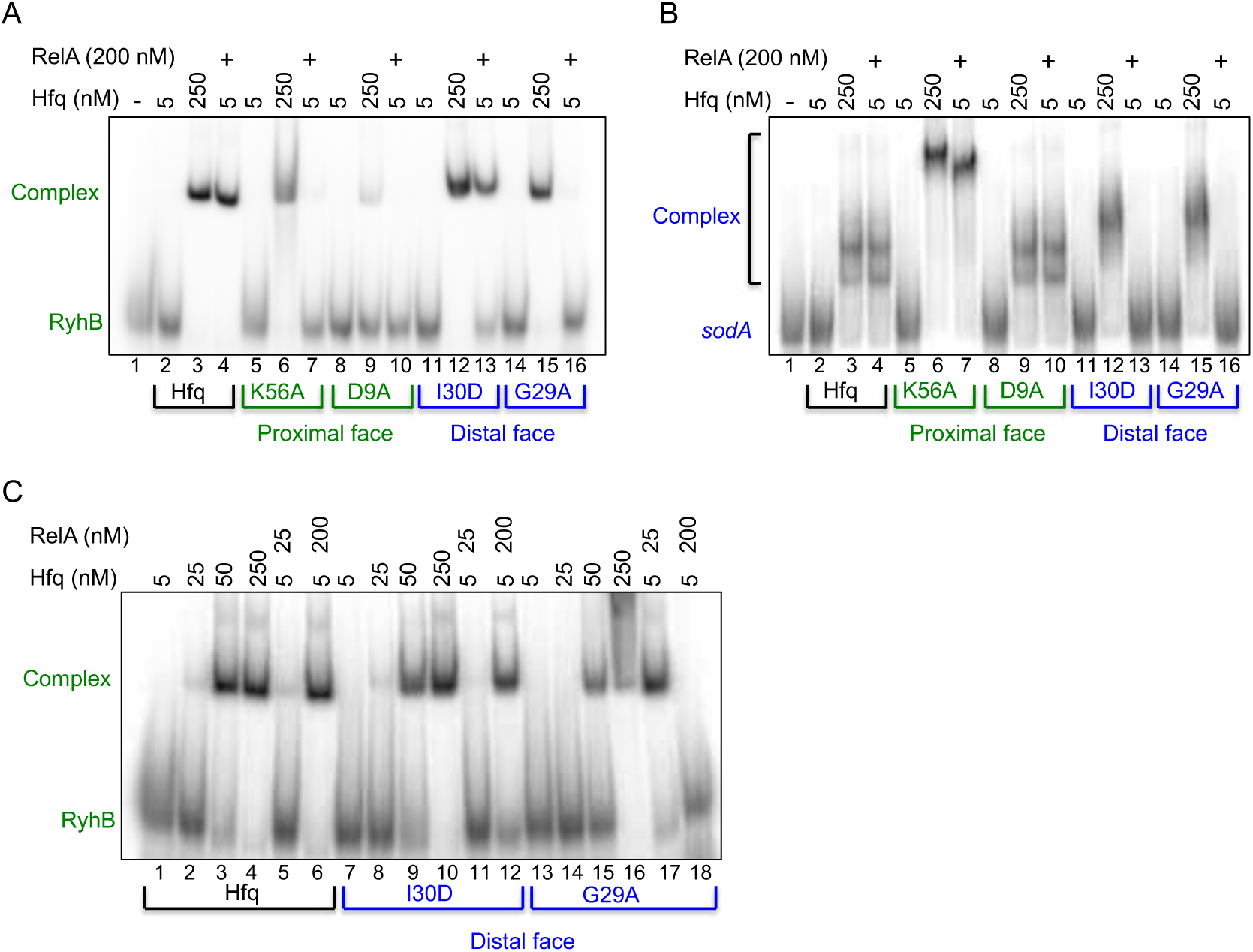
RelA facilitates binding of proximal RNA by Hfq distal mutant and vice versa (EMSA). Wild-type Hfq (black), Hfq proximal mutants K56A and D9A (green) and Hfq distal mutants I30D and G29A (blue) were incubated at 22°C for 10 min. with or without RelA and either (**A and C**) proximal RyhB RNA (green) or (**B**) distal *sodA* RNA (blue). The products were separated by native gel electrophoresis.

**Figure S10.**
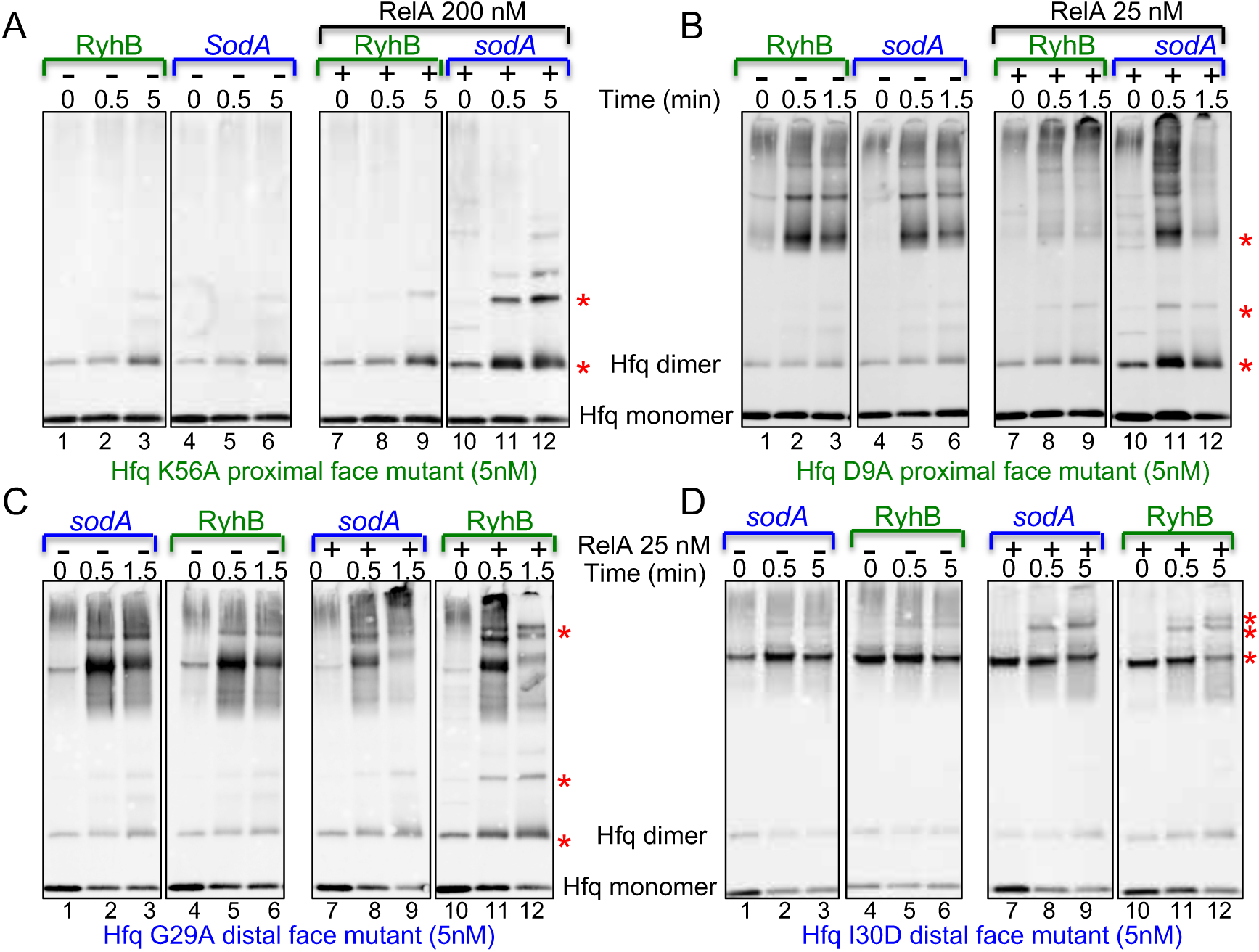
RelA mediated Hfq multimerization requires an initial RNA binding to Hfq (western). Proximal RyhB RNA (green) or distal *sodA* RNA (blue) were incubated at 22°C for 10 min. with proximal Hfq mutants Hfq:K56A (**A**) and Hfq:D9A (**B**) or with Hfq distal mutants Hfq:G29A (**C**) and Hfq:I30D (**D**) and with or without purified RelA protein. Thereafter, the products were UV cross-linked followed by protein crosslinking using 0.2% glutaraldehyde. Samples were collected at the time points indicated and the reactions stopped with 200mM of fresh glycine. The proteins separated in 4-20% MOPS gradient gels were detected using a Hfq. Asterisk denotes the formation of new Hfq multimers. Note that Hfq distal mutants form these multimers in the presence of RelA and RyhB, whereas Hfq proximal mutants form the multimers in the presence of RelA and *sodA* RNA.

**Figure S11.**
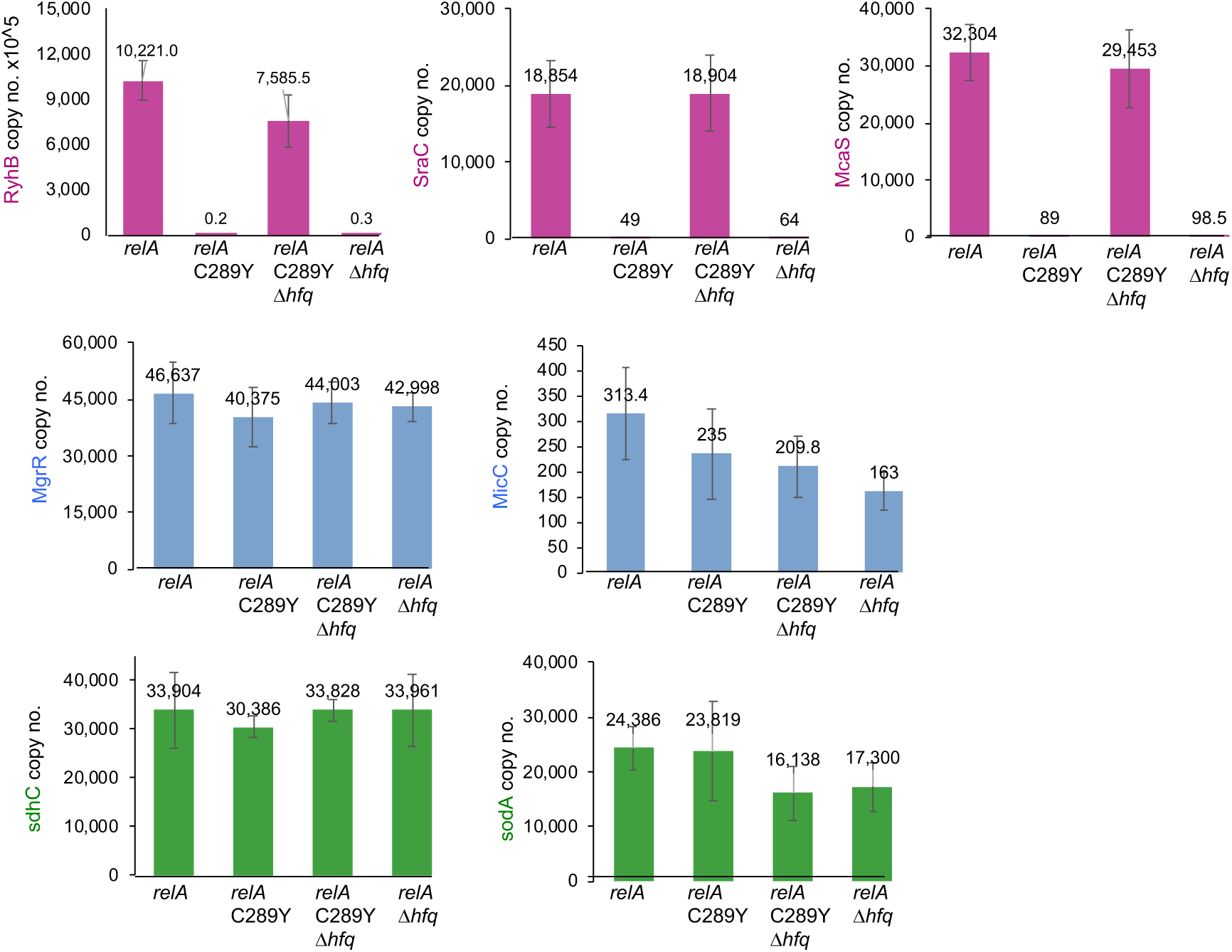
RelA binds sRNAs with GGAG. qRT-PCR of RNA purified during Co-IP from cell lysates as indicated. Two samples of 2-3 biological samples were examined as described in material and methods. sRNAs with GGAG (purple); sRNAs without GGAG (blue); mRNAs with GGAG (green).

**Figure S12.**
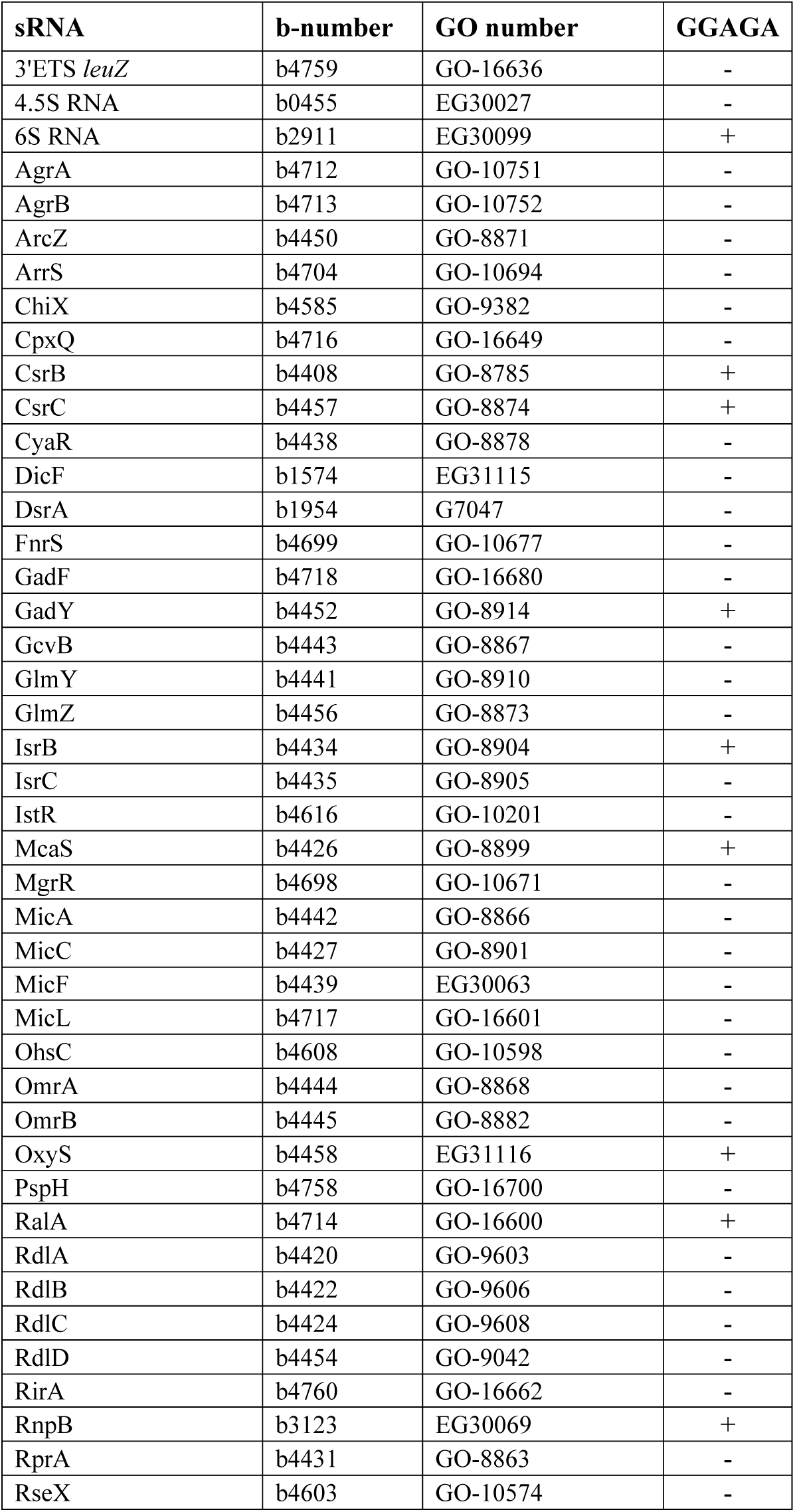
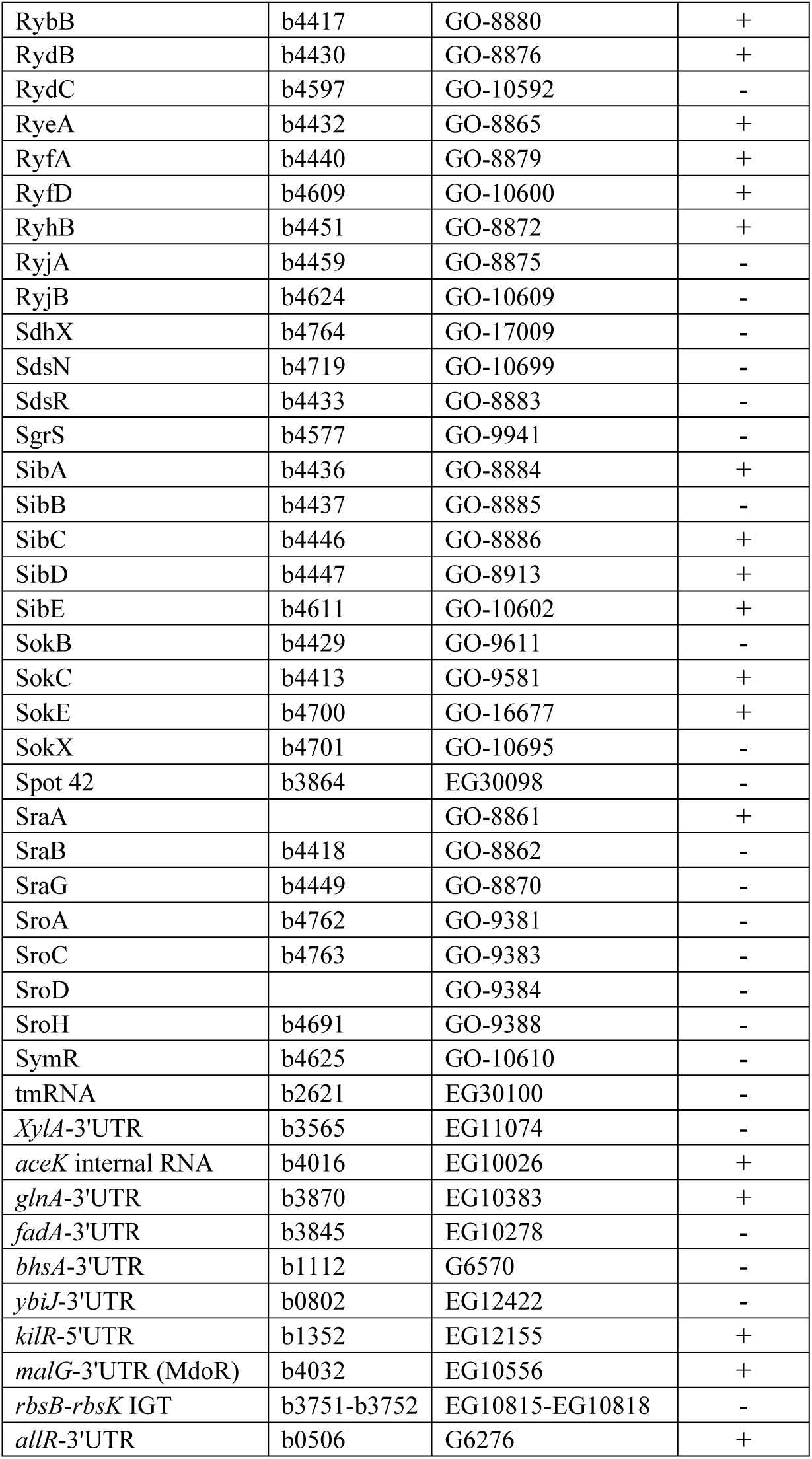
List of sRNAs (*13, 49*) examined for the presence of GGAG sequence.

**Figure S13.**
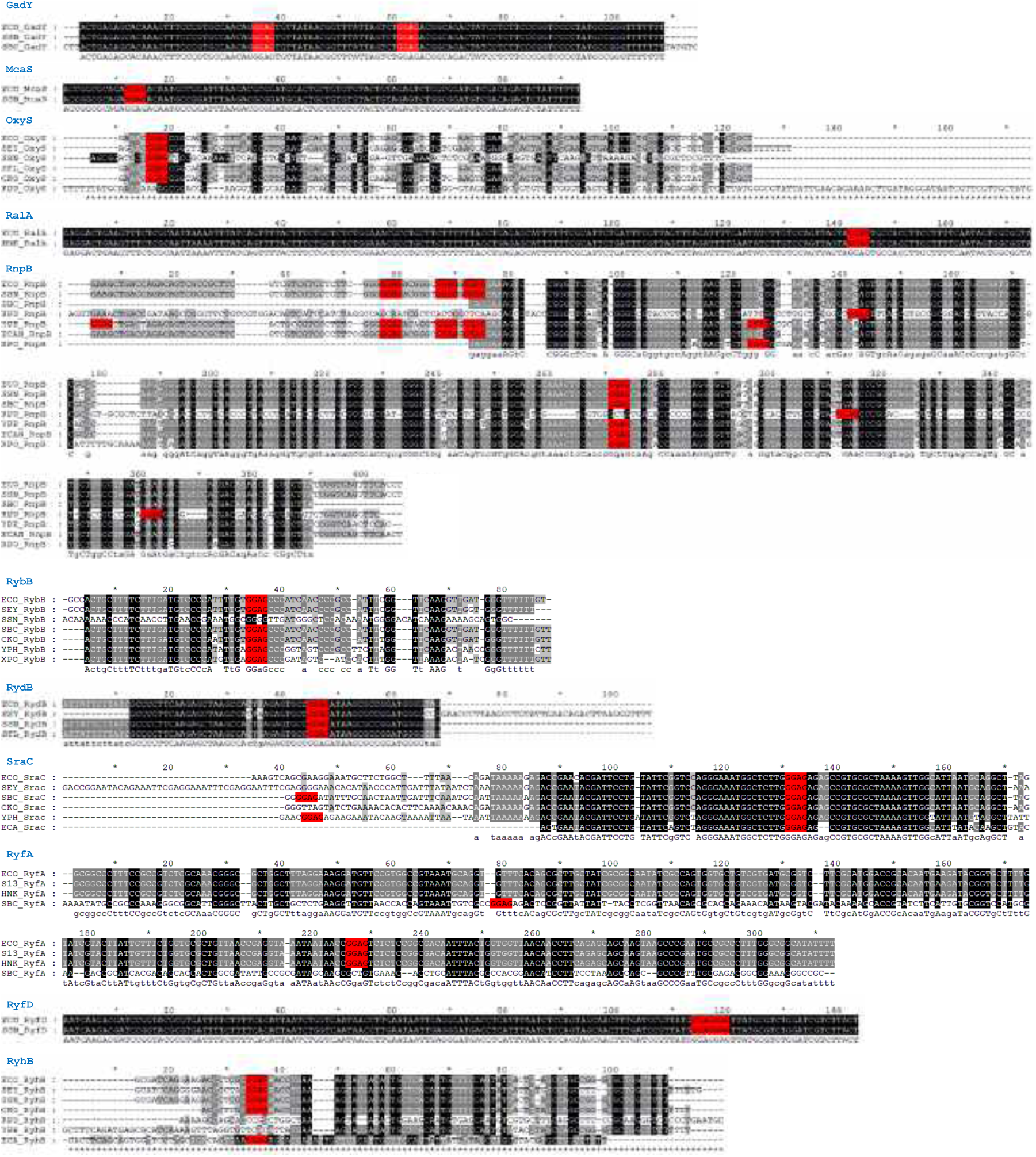
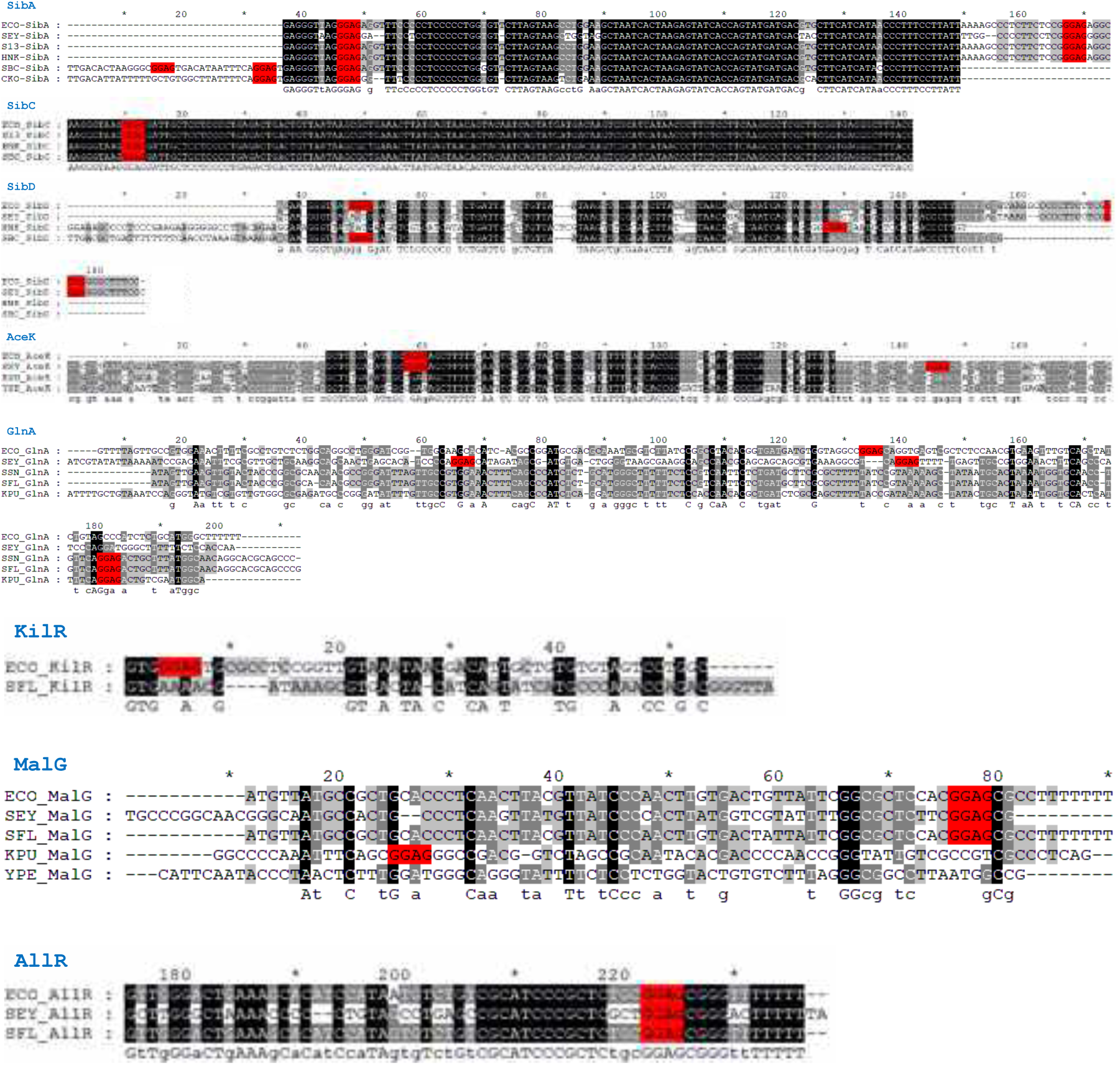
(**A**) Conservation of the GGAG sequence among the sRNAs of diverse enterobacterial species [ECO: *Escherichia coli* K12 MG1655; SEY: *Salmonella enterica subsp. Enterica serovar Typhimurium* SL1344; S13: *Salmonella sp*. S13; HNK: *Salmonella sp. HNK130*; SSN: *Shigella sonnei*; SBC: *Shigella boydii*; SFL: *Shigella flexneri*; CKO: *Citrobacter koseri*; KPU: *Klebsiella pneumoniae subsp. Pneumoniae* NTUH-K2044 (serotype K1); ECA: *Pectobacterium atrosepticum* SCRI1043; ECAN: *Enterobacter cancerogenus*; YPH: *Yersinia pestis Harbin* 35; YPE: *Yersinia pestis* CO92 (biovar Orientalis) and XPO: *Xenorhabdus poinarii*]. Nucleotides in the black regions indicate complete conservation (also indicated by uppercase letters at the bottom of the sequences) while nucleotides in the grey regions indicate partial conservation (also indicated by lowercase letters at the bottom of the sequences). The GGAG sequence is highlighted in red.

**Figure S13.**
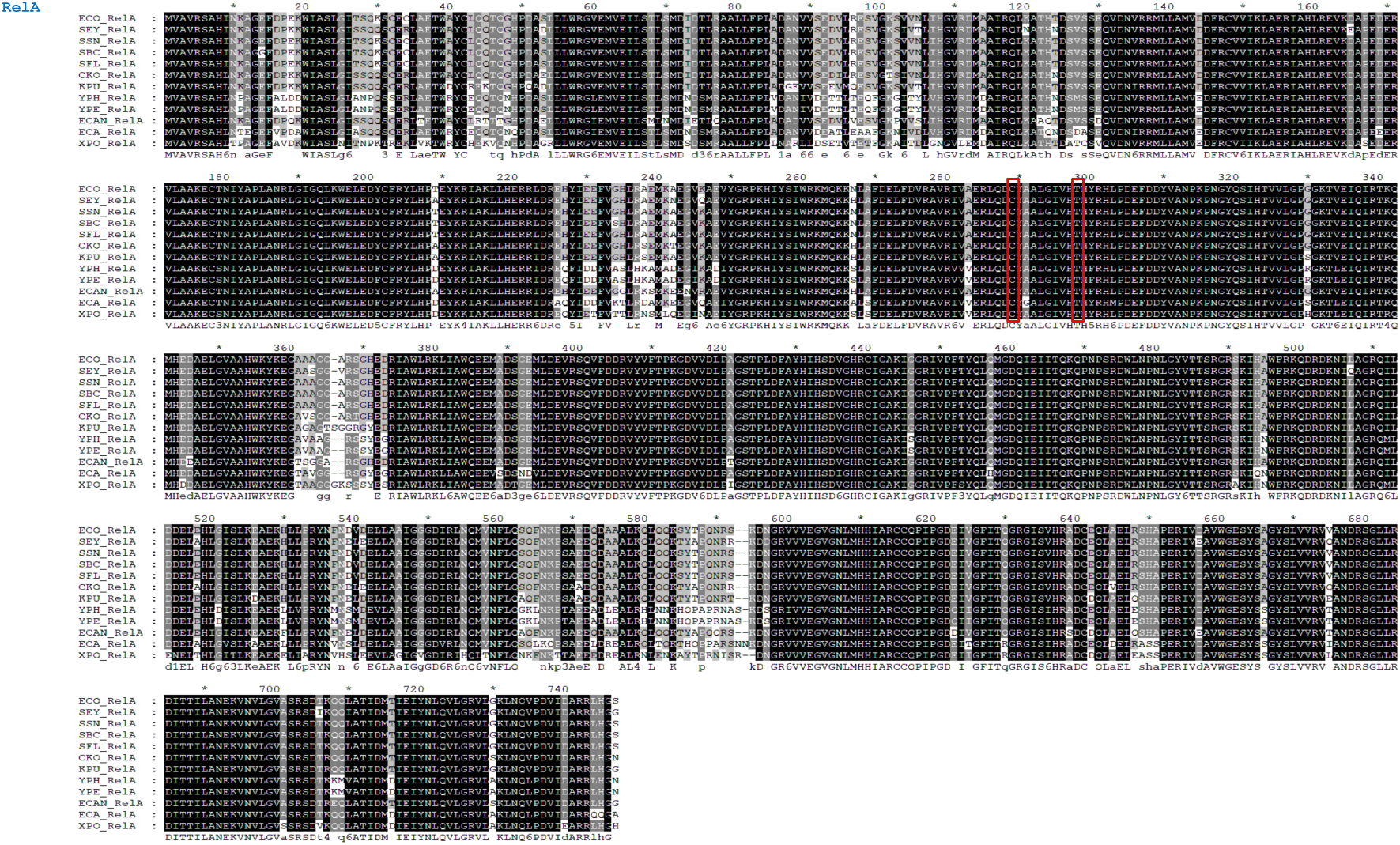
(**B**) Conservation of RelA amino acid sequence among diverse enterobacterial species (see next slide) [ECO: *Escherichia coli* K12 MG1655; SEY: *Salmonella enterica subsp. Enterica serovar Typhimurium* SL1344; S13: *Salmonella sp*. S13; HNK: *Salmonella sp. HNK130*; SSN: *Shigella sonnei*; SBC: *Shigella boydii*; SFL: *Shigella flexneri*; CKO: *Citrobacter koseri*; KPU: *Klebsiella pneumoniae subsp. Pneumoniae* NTUH-K2044 (serotype K1); ECA: *Pectobacterium atrosepticum* SCRI1043; ECAN: *Enterobacter cancerogenus*; YPH: *Yersinia pestis Harbin* 35; YPE: *Yersinia pestis* CO92 (biovar Orientalis) and XPO: *Xenorhabdus poinarii*]. Nucleotides in the black regions indicate complete conservation (also indicated by uppercase letters at the bottom of the sequences) while nucleotides in the grey regions indicate partial conservation (also indicated by lowercase letters at the bottom of the sequences). The amino acids at position 289 (C289) and 298 (T298) are marked by red boxes.

